# Multiple freeze-thaw cycles lead to a loss of consistency in poly(A)-enriched RNA sequencing

**DOI:** 10.1101/2020.04.01.020792

**Authors:** Benjamin P. Kellman, Hratch M. Baghdassarian, Tiziano Pramparo, Isaac Shamie, Vahid Gazestani, Arjana Begzati, Shengzhong Li, Srinivasa Nalabolu, Sarah Murray, Linda Lopez, Karen Pierce, Eric Courchesne, Nathan E. Lewis

**Affiliations:** Department of Pediatrics, University of California, San Diego; Bioinformatics and Systems Biology Program, University of California San Diego; Autism Center of Excellence, Department of Neuroscience, University of California San Diego; Department of Medicine, University of California San Diego; Department of Pathology, University of California San Diego; Department of Bioengineering, University of California San Diego; Novo Nordisk Foundation Center for Biosustainability, University of California San Diego

**Keywords:** RNA-Seq, quality control, freeze-thaw, sample preparation, differential expression

## Abstract

RNA-Seq is ubiquitous, but depending on the study, sub-optimal sample handling may be required, resulting in repeated freeze-thaw cycles. However, little is known about how each cycle impacts downstream analyses, due to a lack of study and known limitations in common RNA quality metrics, e.g., RIN, at quantifying RNA degradation following repeated freeze-thaws. Here we quantify the impact of repeated freeze-thaw on the reliability of downstream RNA-Seq analysis. To do so, we developed a method to estimate the relative noise between technical replicates independently of RIN. Using this approach we inferred the effect of both RIN and the number of freeze-thaw cycles on sample noise. We find that RIN is unable to fully account for the change in sample noise due to freeze-thaw cycles. Additionally, freeze-thaw is detrimental to sample quality and differential expression (DE) reproducibility, approaching zero after three cycles for poly(A)-enriched samples, wherein the inherent 3’ bias in read coverage is more exacerbated by freeze-thaw cycles, while ribosome-depleted samples are less affected by freeze-thaws. The use of poly(A)-enrichment for RNA sequencing is pervasive in library preparation of frozen tissue, and thus, it is important during experimental design and data analysis to consider the impact of repeated freeze-thaw cycles on reproducibility.

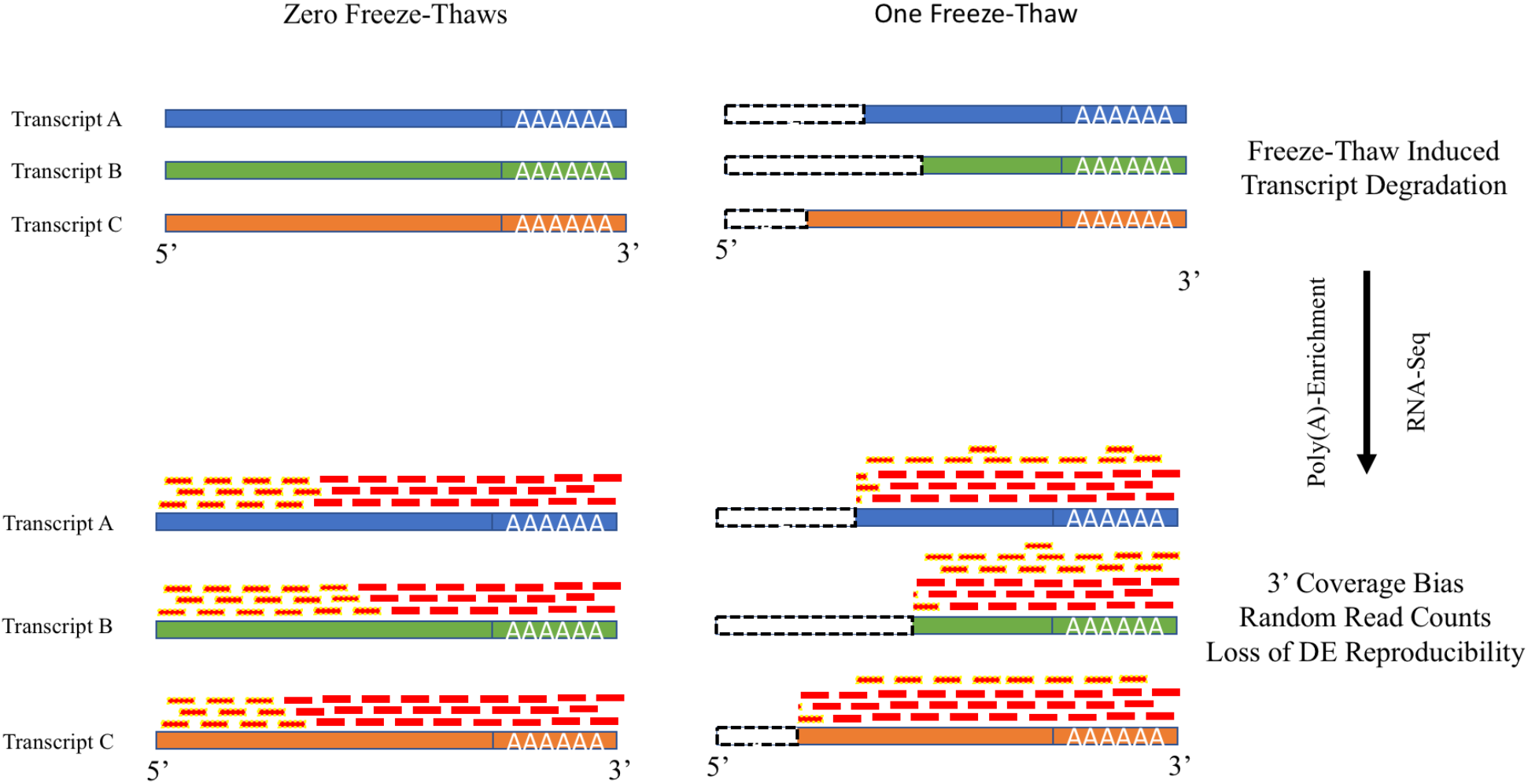

## Introduction

RNA sequencing (RNA-Seq) is a ubiquitous technology, used to answer a wide range of biological questions. Methods for aligning, quantifying, normalizing and analyzing expression data are available through popular packages such as Tophat, STAR, cufflinks, SVA, RUV, Combat, DESeq2, edgeR, Kallisto, Salmon, BWA-MEM, and many others^1–11^. Each method aims to accommodate and mitigate the unique challenges presented by RNA-Seq data. Some approaches attempt to account for characterized variability in RNA-Seq measurements due to factors such as sequencing depth, gene length, and transcripts’ physical characteristics (e.g., GC content). Others account for “unwanted variance” due to technical, batch, or experimental variation. Yet, the influence of sample processing, such as tissue lysis and processing time^12–14^, is not sufficiently characterized such that it can be explicitly controlled. It is important to adequately characterize noise introduced to RNA-Seq measurements by sample processing steps to optimize sample quality, account for transcript degradation, and improve the accuracy and reproducibility of sequencing.

Transcript degradation continues after sample acquisition and affects data quality. Sample storage conditions (e.g. temperature and the use of stabilizing reagents) affect sample quality via RNA degradation^14,15^. Yet, varying sources of degradation can impact RNA-Seq in different manners^16^. Degradation introduces variability in signal and can be impacted by sample handling. Non-uniformity in degradation across genes and samples causes inaccurate normalization and transcript quantification^17^. Poly(A)-enrichment methods are commonly used to separate mRNA from other highly abundant RNA molecules (e.g., rRNA, tRNA, snoRNAs, etc.), but variable degradation directly impacts read counts by causing non-uniform transcript coverage^18^. Of particular interest, freeze-thaw can induce 20% degradation of spike-in standards per cycle, a factor that may be generalizable to mRNA transcripts^19^. Freeze-thaw cycles increase RNA degradation by disrupting lysosomes which store RNases, freeing the enzymes to promiscuously catalyze nuclease activity^20^. Furthermore, partially defrosted crystals create uneven cleaving pressure on mRNA strands^21,22^. Despite these observations, the extent to which freeze-thaw negatively impacts count and differential expression in RNA-Seq analyses has not been comprehensively characterized.

Standard sample quality control often relies on RNA integrity number (RIN), which quantifies the 28S to 18S rRNA ratio^23^. RIN-based quality control approaches rely on a heuristic threshold to assess sufficient quality^24,25^. RIN-based metrics have known confounders such as transcript level, and thus have been called into question as an appropriate quality metric^26^. For example, RIN failed to indicate a decrease in sample quality in lung cancer tissue samples that underwent five freeze-thaw cycles^27^ and, in statistical analyses, failed to correct for the effects of degradation^28^. Despite this, many studies rely on RIN to correct for and assess sample quality confounders^16,29,30^. This is especially problematic in the case of transcript degradation because RIN scores are based on entire samples, while degradation effects can be transcript-specific^17,31,32^. Furthermore, existing studies on degradation are not simply generalizable to freeze-thaw, which has distinct and independent effects on sample quality and must be fully explored as such^16,33^.

Here, we tested the susceptibility of poly(A)-enriched RNA-Seq results after multiple freeze-thaw cycles. We assessed sample quality independently of RIN by simulating read count variability to capture the noise between technical replicates. We found that each additional freeze-thaw cycle increased the random counts between technical replicates by approximately 4%. Subsequently, differential expression reproducibility approached zero after three freeze-thaw cycles. These effects are not captured by RIN. We find that these effects are reflected in increasing 3’ bias in read coverage when combining poly(A)-extraction with freeze-thaw, a phenomena that appears to be generalizable to publically available datasets.

## Results

### 3’ bias in read coverage of public datasets is associated to poly(A)-extraction and freeze-thaw

To examine the prevalence of bias from repeated freeze-thaw cycles, we search for the presence of RNA-seq distortion in publically available datasets. Specifically, we analyze the gene-coverage distribution in samples prepared with either poly(A)-extraction or ribosomal depletion. Since freeze-thaw enhances transcript degradation and poly(A)-enriched samples select mRNA by hybridization to the poly(A)-tail, we expect increased read coverage on the 3’ end of transcripts--3’ bias--when these two factors are combined. To test this expectation, we compared gene body coverage from the 5’ to 3’ end between poly(A)-enrichment and ribosomal depletion prepared samples with and without freezing. Specifically, we examined the median coverage percentile, the percentile-normalized nucleotide at which median cumulative coverage for a given sample is achieved (**Fig. 1a**).

**Figure 1:**
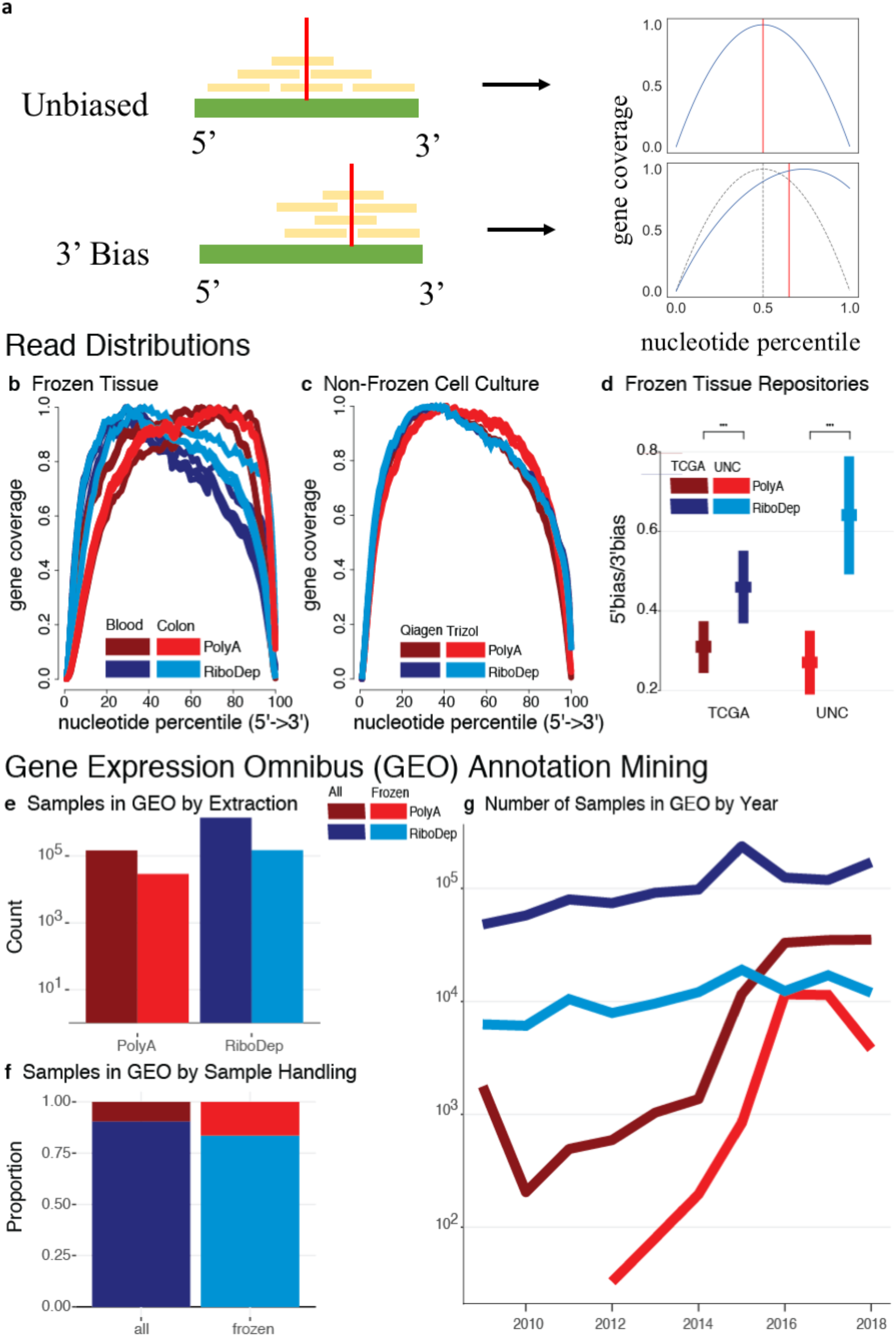
3’ Bias is Exacerbated in Frozen, Poly(A)-extracted Samples Across Multiple Studies: (a) Demonstration for determining median coverage percentile (red vertical line). When coverage is unbiased, reads (yellow) are distributed throughout the entire body of the transcript (green). In the absence of read bias and observing coverage as a function of the nucleotide percentile, we see that cumulative coverage along the transcript reaches 50% half-way through the gene body, at the 50^th^ percentile nucleotide. In contrast, given a 3’ read bias, there is a shift in the distribution of reads and cumulative coverage reaches 50% at, for example, the 60^th^ percentile nucleotide. This results in a rightward shift in median coverage percentile towards the 3’ end of the transcript. In the middle row, gene coverage (y-axis) at the i^th^ nucleotide percentile from 5’ to 3’ (x-axis) is displayed for samples that were extracted using either poly(A)-enrichment or ribosomal depletion. Gene body coverage distributions were calculated for (b) for tissue samples that underwent an unspecified number of freeze-thaw cycles and (c) cell-culture samples samples that underwent no freeze-thaw cycles. (d) Comparison of 5’ to 3’ bias ratio (y-axis) of samples from the TCGA and UNC tissue repositories (x-axis) between extraction methods (two-sample t-test). Quantifying human RNA samples listed in GEO from 2008-2018, and stratifying by those annotated as “frozen”, we observe (d) the number of samples prepared with poly(A)-extraction or ribosomal depletion (x-axis) (gray), (f) the proportion of samples extracted using either method, and (g) the change in the number of samples over time.

We analyzed the gene-coverage distribution in three studies: (1) RNA extraction in previously frozen solid and liquid tissues, (2) RNA extracted immediately after lysis of cultured cells without freezing, and (3) RNA extraction in previously frozen tissue from important public tissue resources.

In the first study^34^, a comparison of the performance of poly(A) and ribosomal depletion in liquid and solid frozen tissue, we see a significant (one-sided Wilcoxon test, p = 0.01591) shift in the median gene coverage percentile towards the 3’ end in the poly(A)-extracted samples (**Fig. 1b**). Using generalized linear regression, we found that the median coverage percentile of poly(A)-extracted samples was 3.88% higher (Wald p = 0.011) than comparable ribosomal depletion extracted samples. In the second study^35^, cells were not frozen before extraction and there was a small and insignificant difference in the 3’ bias associated with library preparation (one-sided Wilcoxon test, p = 0.13, **Fig. 1c**). While the first two studies extract RNA from tissue and cell-culture respectively--tissue extraction is typically lower quality, they are internally controlled and therefore comparable. Finally, the third study^36^ examines the impact of RNA extraction in frozen tissue from the UNC and TCGA tumor tissue repositories. We found a significant (two-sample t-test, p<1e16) decrease in the 5’-to-3’ coverage ratio in poly(A)-extracted samples compared to ribosomal depletion (**Fig. 1d**), indicating an increase in 3’ bias. 3’ bias was consistent across RNA extracted from tissue frozen at either repository.

Next, we explored how widespread the impact of these observations may be by quantifying the prevalence of poly(A)-extraction from frozen tissue by examining metadata in the Gene Expression Omnibus (GEO). With GEOmetadb ^37^, we queried all human RNA samples between 2008 and 2018 using either poly(A)-extraction or ribosomal depletion. There are tens to hundred of thousands of samples annotated as “frozen” for both total and poly(A)-extraction methods (**Fig. 1e**). We note that this is an order of magnitude less than the total query, a value potentially diminished by the complexity of the metadata. In samples annotated as “frozen”, the frequency of poly(A) extraction increases from less than 10% to over 25% **(Fig. 1f)** suggesting that the problematic combination is prevalent and apparently preferred. Finally, stratifying this trend over time, we see that poly(A)-extraction, as well as the relative proportion of poly(A)-extracted frozen samples, is increasing in popularity relative to total RNA extraction, where usage has remained fairly consistent (**Fig. 1g**).Taken together, these results indicate a potential, widespread distortion in RNA-seq associated with a deleterious interaction between poly(A)-extraction and freeze-thaw. To explore this potential more formally, the remainder of our analyses focus on a specific experiment to address this question. Specifically we subjected whole-blood extracted leukocyte samples--with technical replicates--from autistic or typically developing toddlers to a varying number of freeze-thaw cycles, which we record along with other sample quality metrics such as RIN.

### An Additional Freeze-Thaw Cycle Increases Random Read Counts 1.4-Fold

To address the scarcity in quantification of loss in sample quality due to freeze-thaw, we compare changes in sample quality between technical replicates. We first note that neither RIN nor TIN capture significant (one-sided Wilcoxon test) decreases in sample quality due to increased freeze-thaw (**Fig. S1**). Given previous indications that these metrics may not sufficiently address transcript degradation, we instead measure the introduction of noise—the randomness in read counts between technical replicates—to samples by freeze-thaw. We simulated this randomness to reflect the dissimilarity between technical replicates. Since noise does not rely on RIN, we could compare freeze-thaw and RIN-effects independently.

Median noise increased 1.4-fold from one to two freeze-thaw cycles (one-sided Mann-Whitney U test, p ≤ 0.007) on average across all measures (**Fig. 2a**). Noise between technical replicates when samples have only undergone one freeze-thaw was estimated to be 9.11-10.15% (Wald test, p ≤ 5.77e-7). The expected increase in noise per additional freeze-thaw cycle was estimated to be 3.6-4.1 percentage points (Wald test, p ≤ 8.12e-3) (**Fig. 2b**).

**Figure 2:**
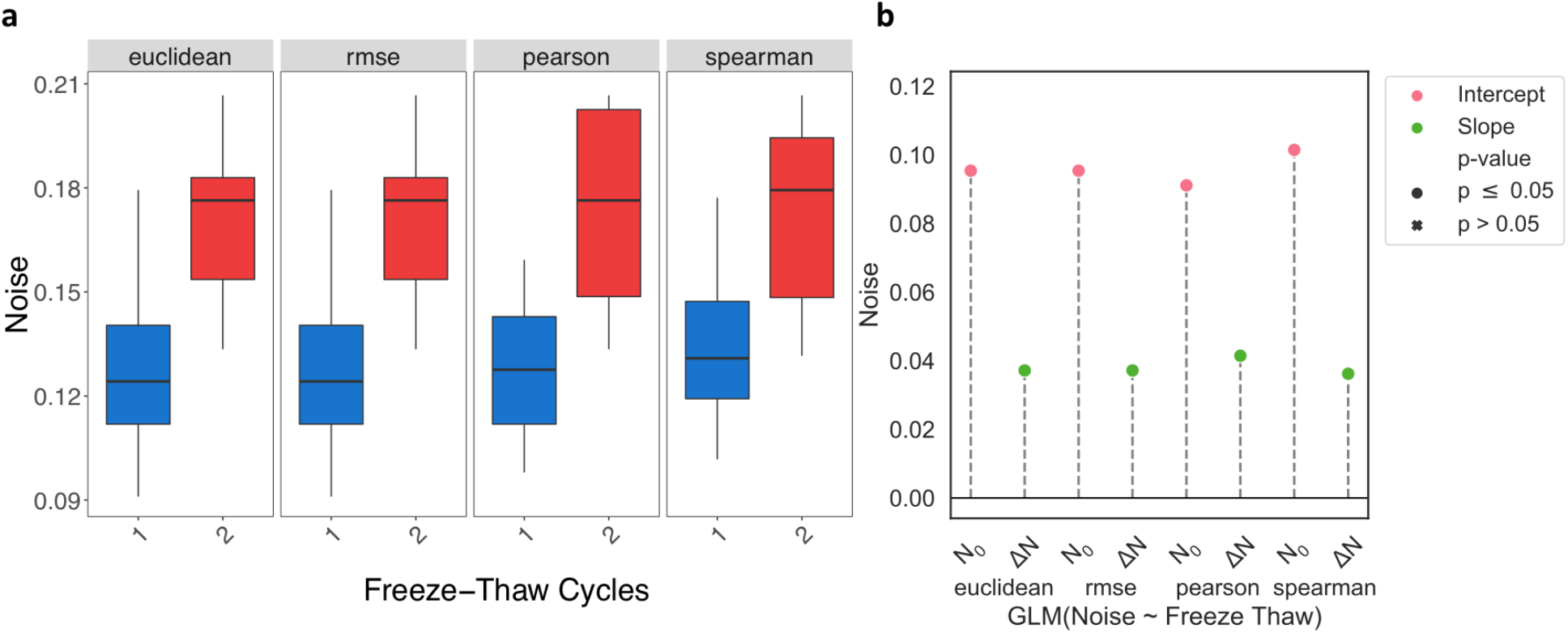
Higher Noise in Samples with More Freeze-Thaw Cycles. From left to right, noise—the randomness in read counts between technical replicates—is estimated using Euclidean distance, RMSE, Pearson correlation, and Spearman correlation. (a) Box plots of noise for samples that underwent either one or two freeze-thaws. (b) A generalized linear model was used to determine the expected noise at one freeze-thaw (N0, pink) and the expected change in noise with each additional freeze-thaw (ΔN, green). All estimates are significant (p ≤ 0.05).

### RIN Does Not Predict Additional Noise After One Freeze-Thaw Cycle

To follow up on our observations that RIN does not sufficiently capture changes in sample quality due to freeze-thaw, we asked whether RIN can reflect the differences in sample quality as measured by noise.

When only considering samples that underwent one freeze-thaw, each unit increase in RIN decreases noise by 3.24-3.38 percentage points for all metrics (Wald test, p ≤ 6.3e-3) (Fig. **3a-b**). Yet, when only accounting for samples that underwent two freeze-thaw cycles, noise does not significantly change as RIN increases. Taken together, these results indicate that while RIN can be a good measure of noise for samples that underwent one freeze-thaw, it does not capture the loss in sample quality induced by two freeze-thaw cycles.

**Figure 3:**
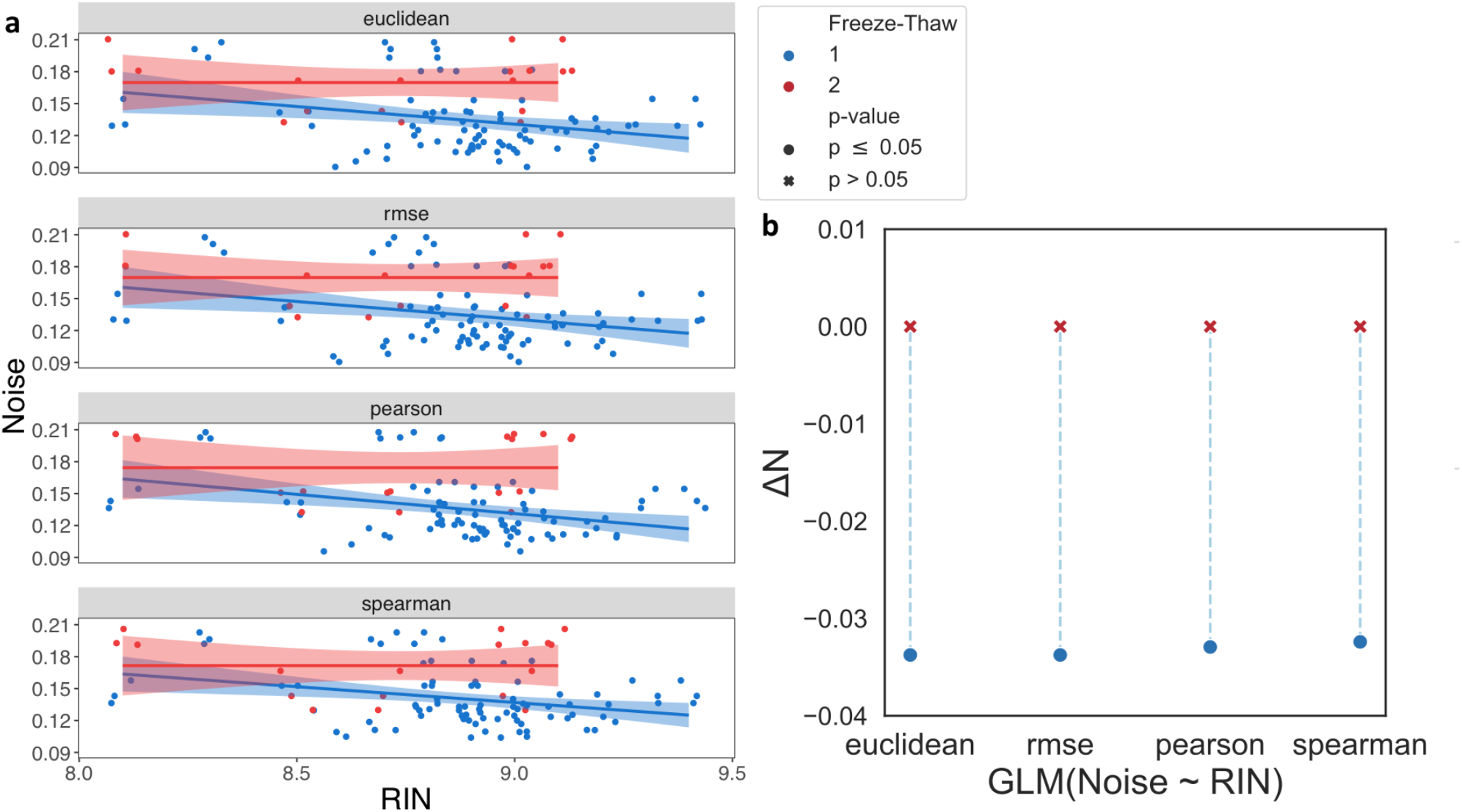
Discrepancy in the Relationship Between Noise and RIN due to additional Freeze-Thaw. Examining the relationship between noise, calculated by Euclidean distance, RMSE, pearson, and spearman correlation, for samples that underwent either one (blue) or two (red) freeze-thaw cycles. (a) Scatter plots comparing noise (y-axis) to RIN (x-axis). The solid lines show a linear regression fit and the shaded regions is the 95% confidence interval for this fit. (b) The expected change in noise due to a one point increase in RIN (ΔN, y-axis) estimated by a generalized linear model. Significant estimates (p ≤ 0.05) are marked by a circle and insignificant estimates are marked by a cross.

### Differential Expression Similarity Increases 10.3% in High Quality Samples

Next, we investigated how the introduction of technical variation, noise, impacts downstream RNA-Seq analysis, specifically, differential expression (DE) analysis. As such, we assessed the reproducibility of DE results on combinations of samples with varying sample quality. We compared DE results between subsets of various sizes (4 - 14 samples). We measure reproducibility using similarity or discordance, based on correlation and dispersion, respectively. Higher similarity and lower discordance each represent higher reproducibility. We use these measures to assess differences that arise between subsets consisting of high quality (low freeze-thaw or high RIN) and low quality (high freeze-thaw or low RIN) samples.

We held two expectations regarding the effect of sample quality on DE reproducibility in the context of similarity: 1) the reproducibility between subsets with high quality samples should be higher than those with low quality samples, and 2) both subset size and sample quality should interact to increase the reproducibility of DE analysis; the increase in stability is reflected by a higher rate of increase in reproducibility with respect to subset size.

As expected, similarity increases with subset size. This is reflected in the upward shift in the similarity distribution with increasing subset size (**Fig. S10**) and the estimated 0.02 (Wald test, p = 2.2e-5) increase in similarity per additional sample (**Fig. 4a**); thus, expected similarity would increase by 0.20 in a subset with 14 samples over a subset with 4 samples. Regression results for each model predicting similarity are reported in **Supplementary Table 4**.

**Figure 4:**
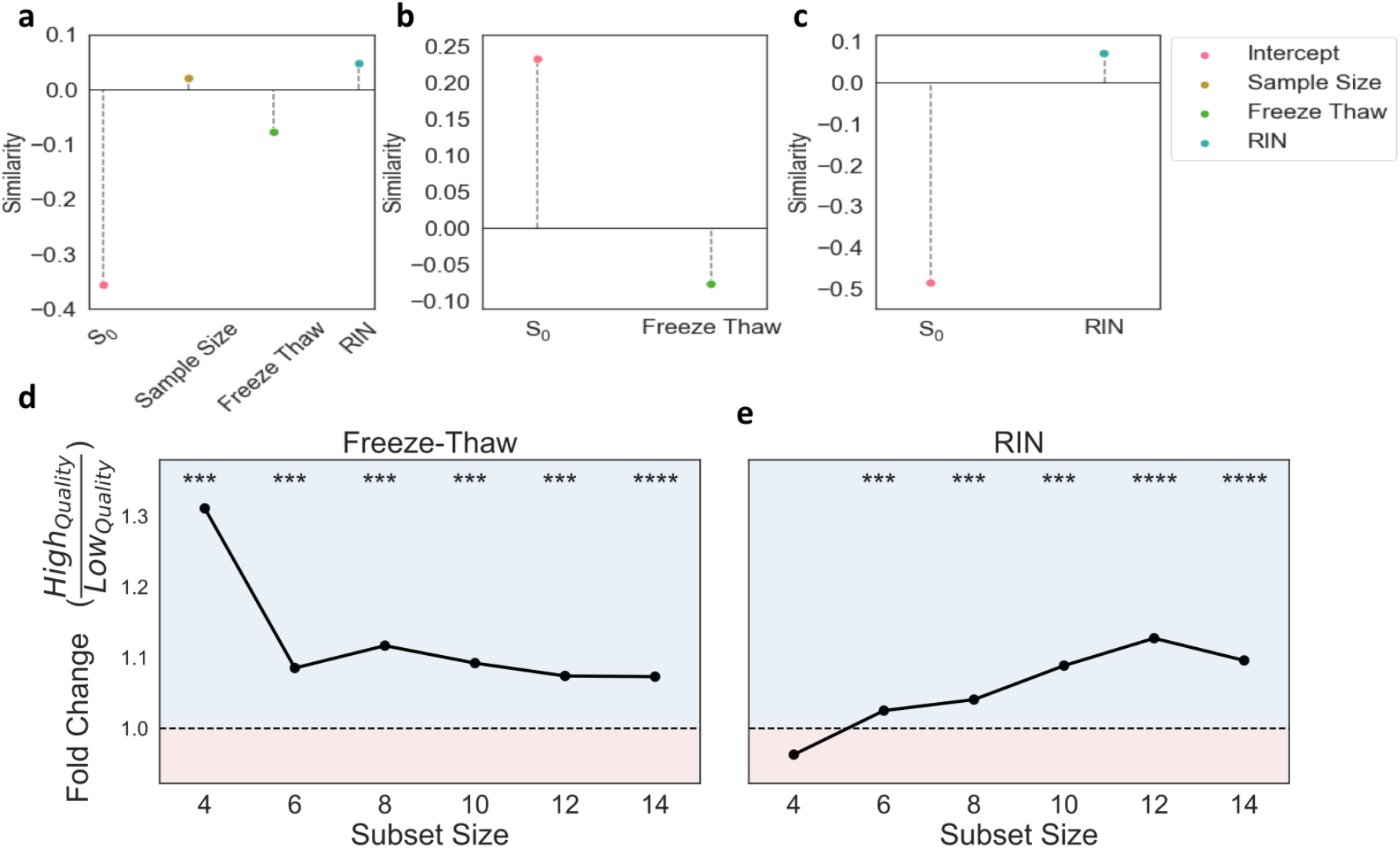
Freeze-Thaw and RIN Both Demonstrate Higher Similarity with Increased Quality. Top panels summarize generalized linear models used to quantify the change in similarity per unit increase in (a) sample size, number of freeze-thaws and RIN combined, (b) only the number of freeze-thaws, and (c) only RIN. S_0_ represents the intercept estimate and sample size, freeze-thaw, and RIN represent coefficient estimates. All estimates are significant. Bottom panels demonstrate fold-change in median similarity of high quality subsets with respect to low quality subsets at each subset size. The region shaded in blue (fold-change > 1) indicates instances where the median similarity for high quality is larger than that of low quality. The region shaded in red (fold-change < 1) indicates instances where the median similarity for low quality is larger than that of high quality. Average (d) freeze-thaw or (e) RIN are used to place subset pairs into high or low quality sample bins. Significance (one-sided Mann-Whitney U test) of comparisons in similarity distributions between high and low quality subset pairs are displayed above each subset size.

We tested our first expectation by placing subset pairs into high and low sample quality bins, defined by either RIN or freeze-thaw, for each subset size and comparing their similarity values. Regardless of sample quality, DE similarity increases with subset size. Yet, for nearly all subset sizes, higher quality bins have significantly (one-sided Mann-Whitney U test, p ≤ 2.8e-17) higher similarity than low quality bins (**Fig. 4e-f**). Across subset sizes, we observed an average 1.13-fold and 1.06-fold increase in similarity from low to high quality samples for freeze-thaw and RIN, respectively.

Similarity significantly (Wald test, p≤ 9.2e-3) decreases with the number of freeze-thaws and increases with RIN when accounting for the effects of sample size (**Fig. 4a-c**), validating our second expectation. Similarity decreases by 0.077 per additional freeze-thaw cycle (Wald test, p = 8.77e-4). Given the estimated similarity of 0.23 for samples that have not undergone freeze-thaw, this implies that DE reproducibility will approach zero after approximately three freeze-thaw cycles (**Fig. 4b)**. Even when accounting for subset size and the effects of RIN, the estimated decrease in similarity from freeze-thaw is nearly the same--0.078 (Wald test, p = 8.77e-4); this further corroborated that RIN alone cannot capture the changes in sample quality due to freeze-thaw. Taken together, these results indicate that higher sample quality increases DE reproducibility as measured by similarity.

### Discordance Decreases ~5-fold In High Quality Samples

We further investigated the relationship between DE reproducibility and sample quality using an effect size sensitive measure of discordance. Specifically, we explored how sample quality affects the relationship between discordance and the mean-variance standardized effect at each subset size. In this context, we expected 1) discordance at any given effect size to be lower in high-quality subsets and 2) the rate of increase in discordance to be lower in high quality subsets relative to low quality subsets.

Corresponding to the regression models used for this analysis, we label our expected discordance when effect size is zero as D0 and the change in discordance per unit increase in effect size as ΔD. We observed a significant (Wald test, p ≤ 9.45e-141) decreasing trend in both D0 and ΔD with increasing subset size (**Fig**. **S11, Supplementary Table 5**).

As expected, independent of sample quality, ΔD demonstrates an overall decreasing trend with respect to subset size for both RIN and freeze-thaw. Given freeze-thaw, at a subset size of 6, there is a 1.1-fold decrease in the value of ΔD from low quality subsets to high quality subsets. The disparity in ΔD between high and low sample quality (Δm = ΔD_Low Quality_ / ΔD_High Quality_) increases nearly monotonically through to the subset size of 14, at which point there is a 3.2-fold decrease; the monotonicity is an indication of the stability of this relation between discordance and sample quality. This causes notable differences in discordance values, even at low effect sizes (**Fig. 5a**).

**Figure 5:**
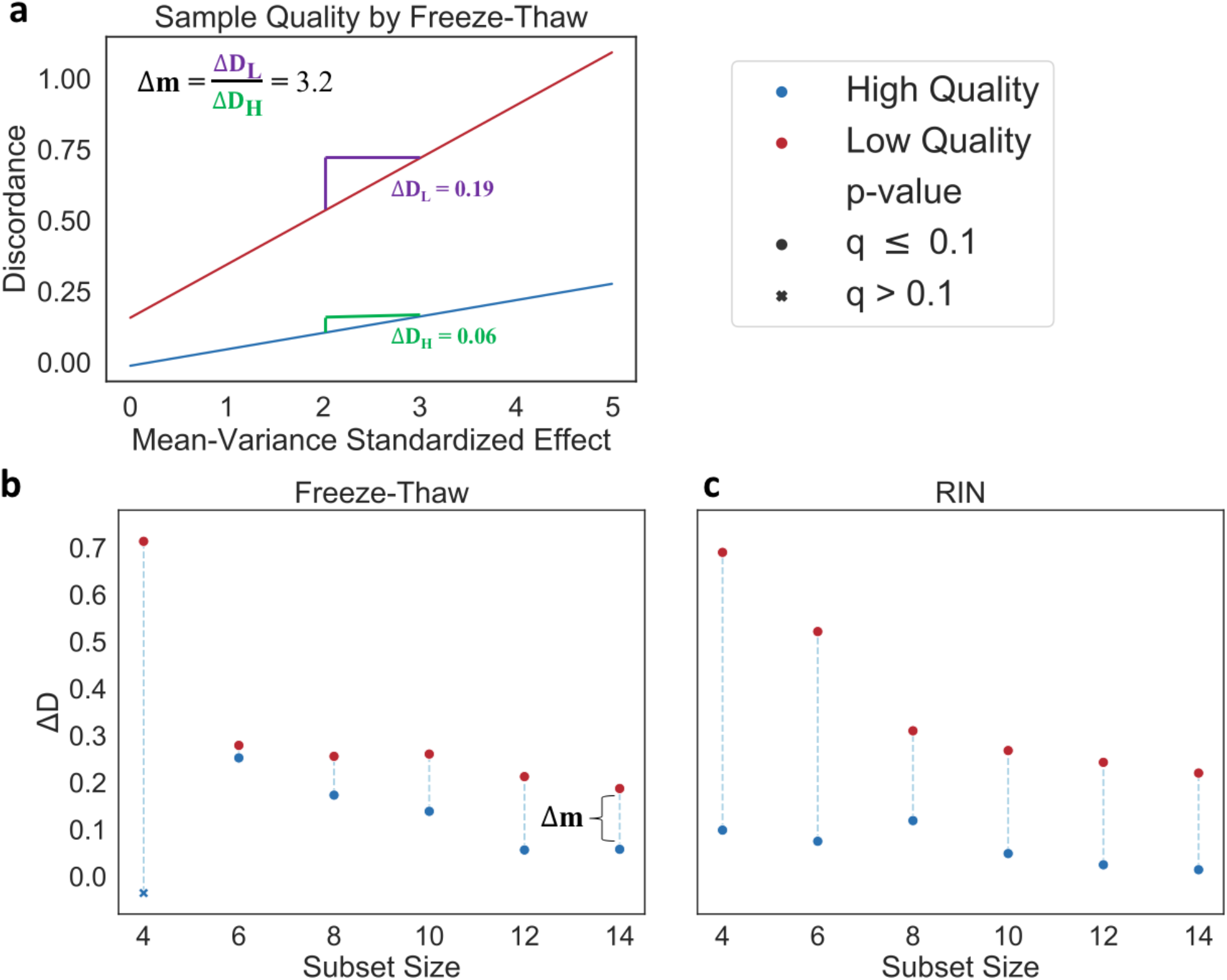
Higher Sample Quality has Lower Discordance at Each Subset Size. GLM estimates of discordance predicted from effect size and subset size at high and low quality subsets at each subset size. High sample quality (blue) is compared to low sample quality (red). ΔD values represent the change in discordance per unit increase in effect size. (a) The predicted discordance with respect to the mean-variance standardized effect at a subset size of 14; sample quality is assessed by freeze-thaw. The disparity (△m) between the change in discordance per unit increase in effect size for high (ΔD_H_) and low (ΔD_H_) quality subsets is also displayed. Summary of results for each subset size (x-axis) for sample quality represented by either (b) freeze-thaw or (c) RIN. Significant estimates (Wald test, Benjamini-Hochberg FDR correction, q ≤ 0.1) are marked by a circle and insignificant estimates are marked by a cross. For freeze-thaw, Δm corresponding to panel A is also displayed.

We estimate discordance with respect to effect size at each subset size and for subsets of either high and low quality (**Fig. 5a**). Consistent with our expectations, D0 and ΔD are consistently lower for high quality subsets as compared to low quality subsets for both freeze-thaw and RIN across all subset sizes (**Fig. 5b-c**). Nearly all estimates are significant after multiple test correction (Wald test, Benjamini-Hochberg FDR correction, q≤ 0.07), with the exception of those for the smallest subset size for freeze-thaw.

Taken together, these results indicate that higher sample quality increases DE reproducibility as measured by discordance.

### Additional freeze-thaw cycles show increased 3’ bias in poly(A)-enriched but not ribosomal depletion samples

Finally, we asked whether repeated freeze-thaw cycles can induce a 3’ bias, consistent with the induction of random reads and the loss of DE reproducibility as well as our initial observation in the public datasets.

Using the median coverage percentile, we found a shift in mRNA coverage towards the 3’ end in the poly(A)-enriched samples relative to ribosomal depletion (**Fig. S13a**). Specifically, the median coverage percentile for poly(A)-enriched samples is significantly (one-sided Wilcoxon test, p < 2.2e-16) larger than that of ribosome depletion (**Fig. S13b**). Samples prepared with poly(A)-extraction have more 3’ coverage bias compared to ribosomal depletion in both one (one-sided Wilcoxon test, p = 3.5e-15) and two (one-sided Wilcoxon test, p = 1.4e-4) freeze-thaw cycles **(Fig. S13d)**. Altogether, this indicates an overall 3’ bias of poly(A)-enriched samples, even independently of freeze-thaw **(Fig. S13b**).

Crucially, this 3’ bias is accentuated when samples are stratified by the number of freeze-thaw cycles (**Fig. 6a**). We observe a significant increase (Wald test, p = 8.3e-4) in median coverage percentile due to the number of freeze-thaw cycles in poly(A)-enrichment. The increase was not maintained when was isolated using ribosome depletion (Wald test, p = 0.155) **(Fig. 6b, Supplementary Table 7)**. For poly(A)-enriched samples, median coverage percentile increases 3.3 percentage points per root freeze-thaw cycle; the square-root of freeze-thaw was used to stabilize variance. We further demonstrate a dependency of 3’ bias on freeze-thaw cycles by showing that median coverage percentile significantly increases with freeze-thaw in poly(A)-enriched samples (Kruskal-Wallis test, p = 0.01). This 3’ bias is particularly apparent after five freeze-thaw cycles (one-sided Wilcoxon test, p = 2.4e-3). In contrast, we observe that median coverage percentile does not significantly change across freeze-thaw cycle counts in ribosomal depletion (Kruskal-Wallis test, p = 0.084) (**Fig. S13c**).

**Figure 6:**
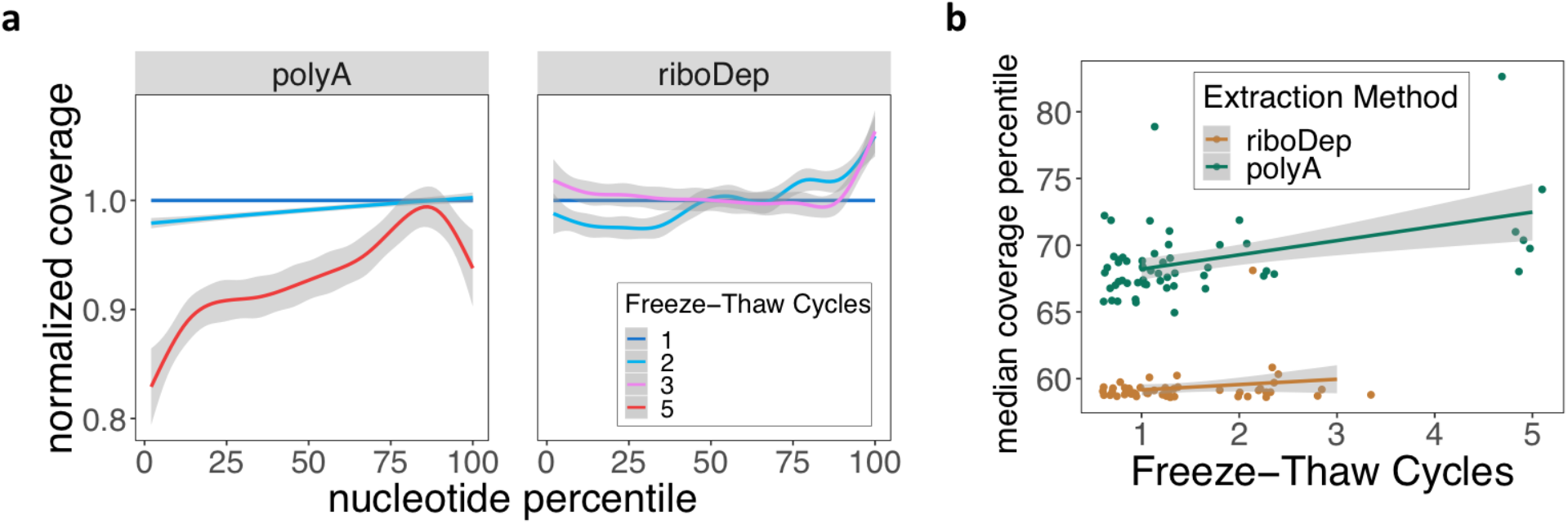
Freeze-Thaw Cycles Exacerbate 3’ Bias in poly(A)-enriched samples. (a) Gene coverage (y-axis) at the i^th^ nucleotide percentile (x-axis) for samples that underwent 1-5 freeze-thaws and were extracted using either poly(A)-enrichment or ribosome depletion. Coverage is normalized to samples that underwent one freeze-thaw. For each sample, coverage is averaged across all genes; samples are aggregated using generalized additive model smoothing, with shaded regions representing 95% confidence intervals. (b) Linear model fits comparing the change in median coverage percentile to the number of freeze-thaw cycles for ribosome depletion (orange) or poly(A)-extraction (green).

Taken together, these analyses indicate that poly(A)-enrichment inherently introduces a 3’ bias in coverage as compared to ribosome depletion, and that this bias is exclusively exacerbated in poly(A)-enriched samples due to freeze-thaw cycles. Thus, 3’ bias may indicate the severity of freeze-thaw induced signal degradation in poly(A)-extracted samples. If this 3’ bias is the root cause of freeze-thaw induced instability in absolute and differential RNA-seq quantification, such instabilities may be subverted by substituting poly(A)-selection for ribosomal depletion during library preparation; such instabilities may be an unnecessary and avoidable inconvenience.

## Discussion

Despite the utility and ubiquity of RNA-Seq, many of the confounding elements associated with the technology are still being characterized. In this work, we demonstrated how one such confounder--sample freeze-thaw--impacts sample quality and downstream analyses. We highlighted biases in publically available datasets, and observed an increased 3’ bias when both freeze-thaw and poly(A)-extraction are features of a sample.

To gain a comprehensive understanding of this effect, we first simulated technical replication to measure the noise between technical replicates with different treatments. This allowed us to examine the impact of freeze-thaw cycles and the ability of RIN to capture those impacts. Next, we examined the impact of freeze-thaw cycles on the robustness and reproducibility of differential expression analysis. We found freeze-thaw cycles were substantially detrimental to the stability of gene expression analysis. By our estimates and at these subset sizes, reproducibility of a differential expression signature approaches zero after three freeze-thaw cycles (**Supplementary Table 4**). Finally, we demonstrated that poly(A)-enriched samples demonstrate substantial 3’ bias in read coverage with increased freeze-thaw cycles. Our results have implications with regards to technical variation due to sample handling, the sensitivity of differential gene expression analysis for frozen tissues and samples, and the utility of RIN.

Technical variation in RNA-Seq is substantial and can be attributed to a variety of factors, including read coverage, mRNA sampling fraction, library preparation batch, GC content, and sample handling^38,39^. As such, accounting for technical variation has been a major research area of focus for the past decade^1,2,5,39,40^. Degradation in combination with poly(A)-enrichment is a known source of variation in RNA-Seq. Yet, before technical variation can be accounted for, it must be characterized. While studies have looked into the effect of degradation on RNA-Seq, each mode of degradation impacts sample quality differently, and direct connections between freeze-thaw and sample quality has mainly been assessed via RIN^16,41,42^.

Our simulation of technical replicates helps delineate technical variation due to sample handling-specifically, freeze-thaw. Furthermore, the resulting noise provides an estimate for the number of random read counts associated with a gene. For example, given an average 25 million reads sequenced per sample, our approximate 4 percentage points difference in noise between one and two freeze-thaw cycles gives an expected stochasticity in 1 million of those reads; approximating the number of protein-coding genes in the human genome to be 20-25 thousand^43^, we can expect a difference of ~40-50 random counts per gene to exist between technical replicates due to a freeze-thaw cycle (**Supplementary Methods, Fig. S15**). Thus, each freeze-thaw introduces a non-negligible level of noise to the quantification of gene expression and differential expression of such genes.

To check for the possibility that there is a signature which can help correct for freeze-thaw distortion of RNA-Seq, we attempt to find a group of common DE genes across various DE methods. We find no such signature (**Supplementary Results**). This is expected, given that a major source of reduced sample quality due to freeze-thaw is mRNA degradation, which occurs randomly for each transcript and sample. A possible path forward is to correct for sample degradation. Several methods have been proposed for this. While some of these methods rely on RIN or similar metrics (e.g. mRIN, TIN, etc.)^16,44^, others have implemented statistical frameworks which account for gene-specific biases. DegNorm, for example, accounts for the gene-specific relative randomness in degradation in its correction approach^17^. Quality surrogate variable analysis (qSVA) specifically improves differential expression by identifying transcript features associated with RNA degradation for its correction^28^. Furthermore, there are recent methods which only assay the 3’ end of a transcript and therefore claim robustness in degraded samples^45^. While these approaches could help account for noise introduced by RNA degradation during repeated freeze-thaw cycles, they cannot necessarily remedy the associated loss of signal.

The effect of freeze-thaw and resultant degradation on RNA-Seq is particularly concerning when considering differential gene expression analysis. It has been observed that RNA degradation can induce the apparent differential expression in as many as 56% of genes^44^. To this end, we quantified this loss of DE reproducibility by measuring similarity and discordance in the context of sample quality. We found a decrease in reproducibility with both decreasing RIN and increasing freeze-thaw. Interestingly, for most reproducibility assessments, we observed a monotonic or near monotonic increase in disparity between low and high quality subsets with respect to subset size. Similarity demonstrated a larger average magnitude of disparity for freeze-thaw, whereas discordance demonstrated a larger average magnitude of disparity for RIN.

Based on our analysis, the utility of RIN in assessing quality when samples undergo freeze-thaw is questionable. The non-uniformity in mRNA degradation^46–49^ due to freeze-thaw sheds light on these challenges, since RIN cannot quantify quality at the individual gene level^23^. This is reflected in the fact that samples with RIN > 8 demonstrate degradation^32^. Furthermore, results assessing the effect of freeze-thaw cycles on RIN are inconclusive. While some studies claim RIN can be used to account for degradation effects in RNA-Seq^16^, others suggest it does not sufficiently capture the effects of degradation on sample quality^26,28^. When directly observing the effect of freeze-thaw on RIN, studies have found a negligible effect^12^ or can only detect an effect after numerous cycles^27,50^.

As such, we re-examined the utility of RIN as a measure of sample quality in relation to our noise estimation of random reads per sample^23^. We found that while noise increases with both decreasing RIN and increasing freeze-thaw, RIN may be an insufficient indicator of quality for samples that have undergone two or more freeze-thaws. Given these results, RIN may not always be a good metric to quantify the difference between technical replicates that have undergone variable sample handling’^7,26–28^. We validate noise by confirming that it does not change with input RNA concentration, excepting outliers **(Fig. S12**). Therefore, in the future, noise could be a useful supplement to RIN when technical replicates are present.

The fact that our predicted decrease in similarity due to freeze-thaw does not change when incorporating RIN into our model further indicates that RIN alone cannot capture the changes in sample quality due to freeze-thaw. Despite this, RIN is a good indicator of sample quality, if not specifically for freeze-thaw. This is reflected in the fact that RIN validates our expectations for DE reproducibility analysis and the comparable range of noise, similarity, and discordance values between freeze-thaw and RIN assessments.

Finally, to confirm our expectation that freeze-thaw decreases sample quality^18–22^ and to further characterize the underlying mechanism, we validated the presence of a 3’ bias in coverage. This builds on our and others’ observations that a lower percentage of poly(A)-enriched transcripts are covered^42^. We compared coverage to ribosome depleted RNA-Seq data, which does not use 3’ hybridization to retain transcripts. We find that poly(A)-enrichment does in fact introduce a strong 3’ bias in coverage as compared to ribosome depletion. This bias is further exacerbated with additional freeze-thaw cycles in poly(A)-enriched but not ribosome depleted samples. This implies that degradation due to freeze-thaw does not impact RNA-sequencing of ribosome depleted samples to the extent that it does in poly(A)-enriched samples. In light of our demonstrations that 3’ bias is associated with a substantial increase in noise and a decrease in DE reproducibility, these findings suggest that RNA-seq from samples that have both been poly(A)-extracted and undergone freeze-thaw cycles likely has unknown, diminished stability. While not all studies have technical replicates to estimate noise, the presence of exaggerated 3’ bias when poly(A)-extraction is combined with freeze-thaw can be a simple indicator of RNA-seq distortion.

## Conclusion

Altogether, these results indicate that transcriptomics quality control steps cannot rely on RIN alone for samples that have undergone poly(A)-enrichment and multiple freeze-thaws. Furthermore, accounting for the effect of freeze-thaw on poly(A)-enriched RNA sequencing is crucial. poly(A)-enrichment is prevalent for RNA-sequencing, and, in parallel, samples that undergo multiple freeze-thaws are common in many protocols, especially rare tissues, e.g., postmortem neural tissue. Yet, there is no clear recommendation to avoid poly(A)-enrichment following multiple freeze-thaws. Together, these results indicate that ribosomal depletion could be a better alternative when freeze-thaw is necessary.

## Declarations

### Ethics Approval and Consent to Participate

In this study, we performed transcriptomics analyses of blood samples drawn from male toddlers with the age range of 1–4 years. Research procedures were approved by the Institutional Review Board of the University of California, San Diego. Parents of toddlers underwent informed consent procedures with a psychologist or study coordinator at the time of their child’s enrollment. Additional details for the recruitment protocol are executed as described in Gazestani et al.^51^

### Author contributions

B.P.K., N.E.L., T.P., S.M., and E.C. designed and planned the experiments. T.P., S.N., K.P., S.M. and L.L. collected the samples, managed diagnostics, conducted transcriptome assays and managed the data. B.P.K, H.M.B. and V.G. planned and conducted analyses. I.S. performed RNA-seq QC. A.B. performed functional enrichment analyses. S.L. wrote the RNA-Seq processing pipeline. B.P.K., H.M.B, and N.E.L wrote the manuscript. N.E.L. supervised the project.

## Acknowledgements and Funding

This work was supported by NIMH R01-MH110558 (E.C., N.E.L., T.P., V.G., S.N., K.P.,L.L.), NIDCD R01-DC016385 (E.C., T.P.), T32GM008806 (H.M.B.), R35 GM119850 (B.P.K., I.S.), the Simons Foundation (E.C.), and generous funding from the Novo Nordisk Foundation through Center for Biosustainability at the Technical University of Denmark (NNF10CC1016517 S.L.).

## Availability of Data and Materials

The datasets supporting the conclusions of this article are available in the NCBI Sequence Read Archive repository (PRJNA627540, https://www.ncbi.nlm.nih.gov/Traces/study/?acc=PRJNA627540&o=acc_s%3Aa). Associated metadata is available in Supplementary Tables 1-2.

## Methods

### Sample Collection and Storage

Blood samples drawn from male toddlers with the age range of 1-4 years were usually taken at the end of the clinical evaluation sessions. To monitor health status, the temperature of each toddler was monitored using an ear digital thermometer immediately preceding the blood draw. The blood draw was scheduled for a different day when the temperature was higher than 37 °C. Moreover, blood draw was not taken if a toddler had some illness (for example, cold or flu), as observed by us or stated by parents. We collected 4–6 ml blood into EDTA-coated tubes from each toddler. Blood leukocytes were captured using LeukoLOCK filters (Ambion). After rinsing the LeukoLOCK filters with PBS, the filters were flushed with RNAlater (Invitrogen) to stabilize RNA within the intact leukocytes. After RNA stabilization, the LeukoLOCK filters were immediately placed in a −20 °C freezer. Additional RNA standards were sourced from normal human peripheral leukocytes pooled from 39 Asian individuals, ages 18 to 47 (Takara/ClonTech: 636592). The RNA standards underwent 1-5 simulated freeze-thaw cycles; 24hrs frozen at −80 °C and 1hr defrosting on ice.

### RNA Extraction, Sequencing and Quantification

For 47 samples (from 16 individuals), mRNA was extracted using polyA selection with the TruSeq Stranded mRNA library preparation kit (Illumina). Ribosomal depletion was used to prepare an additional 52 samples. Relevant metadata regarding polyA-enriched and ribosomal depleted samples can be found in **Supplementary Table 1-2**. Ribosome depletion prepared samples used the TruSeq Stranded Total RNA with RiboZero Gold library preparation kit (Illumina). RNA Integrity Numbers (RIN) were measured using a NanoDrop ND-1000 (ThermoFisher). Poly-A selected samples were sequenced using 50-base pair single end sequencing on a HiSeq4000 (Illumina) to a depth of 25M reads. The ribo-depletion prepared libraries were sequenced using 100-base pair paired end sequencing on a HiSeq4000 (Illumina) to a depth of 50M reads.

Fastq files for each sample underwent quality control using FastQC (v0.33). PolyA and adaptor-trimming were conducted using Trimmomatic ^52^. Reads were aligned to the gencode annotated (v25) human reference genome (GRCh38) using STAR (v2.4.0) ^7^. Alignments were processed to sorted SAM files using SAMtools (v1.7) ^53^. Finally, HTSeq using default settings (v0.6.1) was used to quantify reads ^53,54^.

### Estimation of noise between technical replicates

To estimate the noise between technical replicates of the same individual blood samples, we simulate random loss and gain of reads **(Fig. S2)**. One technical replicate was chosen as the “reference” replicate, making the other technical replicate the “target” replicate. Specifically, we designated replicates that have undergone one freeze-thaw as the reference, and those that underwent two freeze-thaw cycles as the target. The dissimilarity between replicates is measured by one of four metrics (Euclidean distance, RMSE, Pearson correlation, and Spearman correlation). We iteratively add and remove random reads to the reference replicate until the dissimilarity between the simulated replicate and the reference replicate was equal to the dissimilarity between a target replicate and the reference replicate (**Fig. S3, Figure S2**). We define the noise between the reference and target replicate as the fraction of reads added or removed per total reads in the reference replicate to achieve the aforementioned level of dissimilarity. We represent this as a percentage, e.g. 5% noise between a reference and target replicate can be interpreted as 5% randomness between their reads. For additional details on noise simulation, see Supplementary Methods.

### Measuring the Effect of Sample Quality on Noise

To measure the association between noise and sample quality metrics (number of freeze-thaw cycles, input RNA concentrations, and RNA integrity number), we used a generalized linear model (GLM). The significance of the model parameters is determined by the Wald test. All results are reported in **Supplementary Table 3**.

For each model, to mitigate the contribution of potential confounding variables, samples with input RNA concentrations in the top and bottom 5% (|z| ≥ 1.645) were removed, decreasing the total number of samples from 47 to 41. For noise prediction from concentration, samples with more than one freeze-thaw were also excluded, decreasing the total number of samples to 35. Noise prediction from RIN was run separately for samples that had undergone one freeze-thaw and samples that had undergone two freeze-thaws.

### Differential Expression Analysis

We assess whether the observed sample qualities (freeze-thaw and RIN) have an impact on differential expression (DE) reproducibility using a bootstrapping approach. DE was run on random sample subsets of varying sizes (**Fig. S4**). Before subsetting, we filtered our expression matrix for genes with an average count ≤ 20 across all samples. This reduced the number of genes from 10,028 to 4,520. The total number of samples considered was 46 when disregarding samples that were industry standards, were not assigned to either an ASD or TD indication, or did not have a recorded sample quality (RIN or freeze-thaw) value.

We generated subsets containing N = 4-14 samples. For each subset size N, we generated 2,000 unique subsets. Each subset had an equal number of TD or ASD samples. Additionally, only one replicate from each blood sample could be included. These requirements limited our subset size to a maximum of 14 samples.

DE between ASD and TD subjects was conducted using DESeq2^1^. **Fig. S10** summarizes DE results for all subsets. To account for potential confounders, we used RUV^5^. Specifically, we use a set of “in-silico empirical” negative control genes, including all but the top 5,000 differentially expressed genes as described in section 2.4 of the documentation for RUVseq (http://bioconductor.org/packages/release/bioc/vignettes/RUVSeq/inst/doc/RUVSeq.pdf). We check that RUV produces consistent results with previous Autism leukocyte gene expression signatures^51,55^, see **Supplementary Results**.

### Similarity to Assess Differential Expression Reproducibility

To assess DE reproducibility, we measure the similarity in log-fold-change (LFC) values between DE runs. Similarity is calculated as the pairwise spearman correlation of LFC between all subsets of the same size **(Fig. S6**). Genes with a median base mean (the mean of counts of all samples, normalizing for sequencing depth) or median LFC in the bottom 10^th^ percentile across all subsets were excluded, filtering for low magnitude effects **(Fig. S5**).

Average RIN and freeze-thaw were measured for all subset pairs. Resulting distributions for all collected values from similarity analyses are displayed in **Fig. S7**.

Next, subsets of each size were split into two quantile bins for each quality metric separately. High sample quality bins (low freeze-thaw or high RIN) were compared to low sample quality bins. High sample quality subsets were tested for higher similarity than low sample quality bins using a one-sided Mann-Whitney U test.

Additionally, three generalized linear models (GLMs) were fit to quantify the contribution of sample quality metrics to the change in similarity for DE results across subsets. We fit one model to predict similarity from freeze-thaw and RIN, while also accounting for the improvement in reproducibility due to increase in subset size (Similarity ~ Freeze-Thaw + RIN + Subset Size). We also fit two models predicting similarity from freeze-thaw or RIN alone.

### Discordance to Assess Differential Expression Reproducibility

We adapted a measure of concordance to measure discordance, or the lack of reproducibility, between differential expression results^56^. Average RIN and freeze-thaw were calculated for each subset **(Fig. S8-9**). Subsets for each subset size were split into two quantile bins for either quality metric (RIN and freeze-thaw). Genes with a median base mean across all subsets in the bottom tenth percentile were excluded from the analysis **(Fig. S5**).

We do not use the original concordance at the top (CAT) metric because we are not comparing our results to a gold standard dataset. Instead, we use gene-wise LFC standard deviation across subsets as a measure of discordance. Thus, the average LFC for each gene across DE runs is analogous to the gold standard, and the dispersion from this average indicates a lack of reproducibility. At each combination of subset size and quality bins, we calculate discordance and compare it to the gene-wise median effect size **(Fig. S8**). We measure effect size as the mean-variance standardized effect^1^. This and two additional effect size metrics (Cohen’s d and absolute median LFC) we use are further described in **Fig. S9**. Results for all three effect size metrics reflect similar trends and can be found in **Supplementary Tables 5-6**.

We used a GLM to predict discordance from effect size at each subset size. Additionally, in a separate GLM, we account for the interaction between effect size and sample quality (Discordance ~ Effect Size x Sample Quality) at each subset size. Here, sample quality is a dummy variable, assuming a value of 0 for low quality and 1 for high quality. We did not include a term for subset size because regressions were fit within each subset size.

### Read Coverage Bias

The distribution of read coverage over each gene body was measured using *geneBody_coverage.py* from the *RSeQC* (v3.0.0) package ^57^. We measure this coverage ranging from the 0^th^ percentile (5’ end) to the 100^th^ percentile (3’ end) nucleotide. The i^th^ percentile nucleotide is calculated as *nucleotide_i_* / *length_gene_*. Coverage at the i^th^ percentile nucleotide is normalized across all genes within a sample.

For a given sample, the median coverage percentile is defined as the nucleotide percentile at which median cumulative coverage is achieved; cumulative coverage is aggregated from the 5’ end to the 3’ end. The larger the median coverage percentile value, the larger the 3’ bias in coverage. We include 9 industry standards to our analysis--six of which had undergone five freeze-thaw cycles and three of which had undergone one freeze-thaw cycle--to explore the impact at higher freeze-thaw counts. We also include ribosomal depletion extracted samples as a negative control.

We extended this read coverage bias to three additional datasets, all of which contained both poly(A)-extracted and ribosomal depletion extracted samples. The first dataset (PRJNA427184) contains samples for liquid and solid frozen tissue ^34^. The second (PRJEB4197) contains HEK293 samples that explicitly never underwent freeze-thaw cycles ^35^. The third (phs000676.v1.p1) contains frozen tissue samples from the UNC and TCG tumor tissue repositories ^36^. For the first two datasets, fastq files were aligned using STAR and gene body coverage was calculated from alignments using RSeQC as previously described. We did not directly analyze the raw files from TCGA or UNC, but instead reanalyzed the reported 5’ to 3’ bias ratios. This ratio is similar to the inverse of the median coverage percentile: the smaller it is, the larger the 3’ bias in read coverage.

