## Supplementary Text and Figures for "Multiple freeze-thaw cycles lead to a loss of consistency in poly(A)-enriched RNA sequencing"

**Supplementary Figures** 4

Figure S1 - Illustration noise generation to simulate technical replication 4

Figure S2 - Illustration of the noise estimation method comparing a reference sample to a technical replicate as noise is simulated 6

Figure S3 - Illustration of the method used to generate the subsets used for differential expression analyses 7

Figure S4 - Aggregate MA plots for subset differential expression analysis 8

Figure S5 - Illustration of similarity calculation between subset differential expression analyses 10

Figure S6 - Similarity, RIN, and freeze-thaw distributions in similarity analyses 11

Figure S7 - RIN and freeze-thaw distributions across various subset sizes for discordance analysis 12

Figure S8 - Method for calculation of discordance 13

Figure S9 - Distribution of differential expression similarities for different subset sizes 14

Figure S10 - Spread of differential expression discordance for different subset sizes 15

Figure S11 - Concentration does not affect noise 16

Figure S12 - 3' Bias in polyA-extracted samples 17

Figure S13 - Distribution of RIN and concentration with respect to noise 18

Figure S14 - Projected random counts for as a function of library size, number of active genes, and percent noise 19

**Supplementary Methods** 20

Simulation of Technical Replicates 20

Freeze Thaw GSEA 21

**Supplementary Results** 21

Changes in Concentration Have a Negligible Effect on Noise 21

Using RUV in DE Recovers an Autism Signal 22

GSEA Does Not Identify an Apparent Freeze-Thaw Gene Signature 22

**Supplementary Discussion** 22

**References** 24

### Supplementary Figures

#### Figure S1 - Comparison of sample quality metrics between extraction methods and freeze-thaw cycles

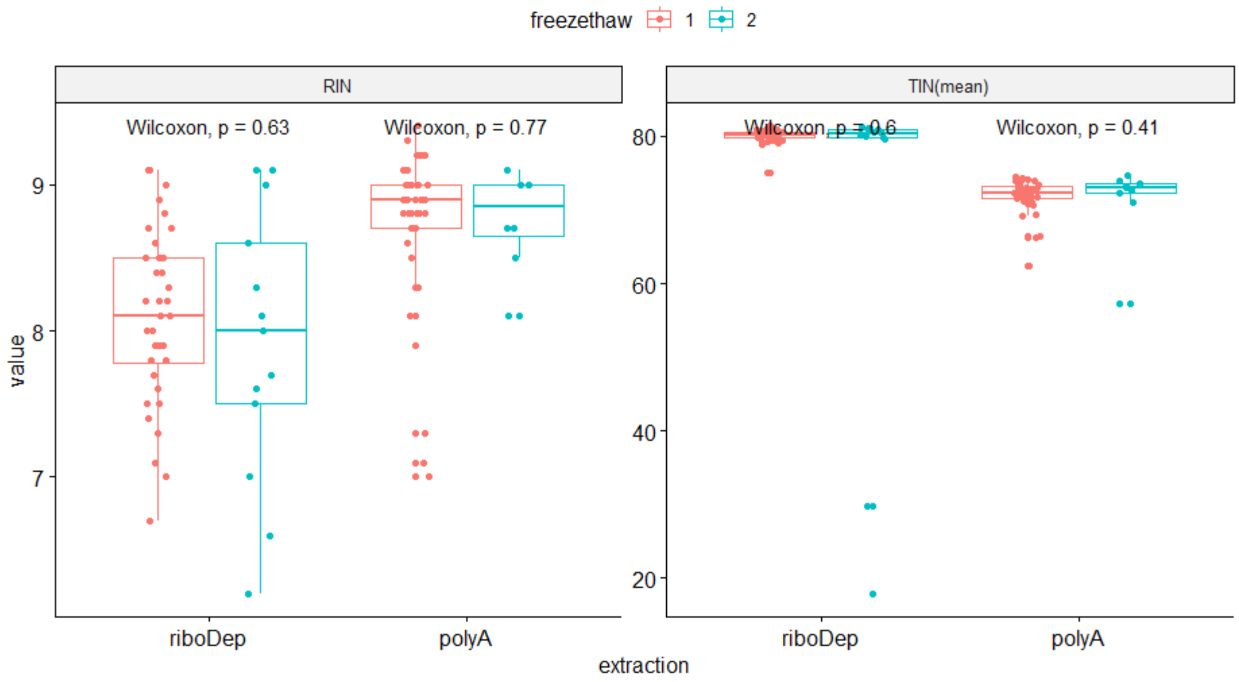


For each extraction method (ribosomal depletion and poly(A)-extraction), we compare RIN (left panel) and TIN (right panel) values between samples that underwent either one freze-thaw (red) or two (freeze-thaws) blue. We find no significant changes in any of the comparisons using a one-sided Wilcoxon test.

#### Figure S2 - Illustration noise generation to simulate technical replication

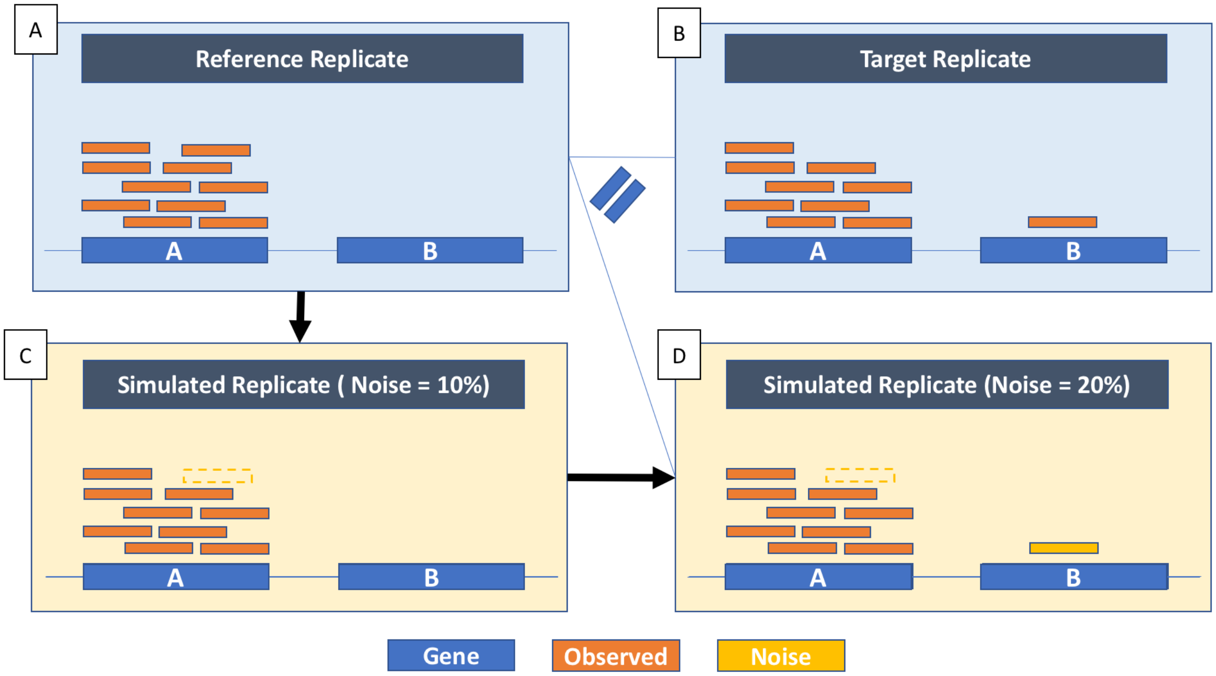


Illustration of noise to simulate technical replication. Poisson simulated technical noise measures noise between technical replicates of the same blood sample. In this toy example, each sample genome has 2 genes. (A) The reference replicate has 10 and 0 reads in gene A and gene B, respectively. (B) In contrast, the target replicate has 9 reads and 1 read in each gene. This results in 20% randomness in reads between the two replicates, which results in differences in both coverage and depth between the two samples. (C) Noise is simulated in the reference replicate to 10% by removal of a read from gene A. (D) Noise is further introduced to the simulated replicated, this time by adding a read to gene B. At this point, the simulated replicate has 20% noise and a dissimilarity from the reference replicate that is equal to the dissimilarity between the target and reference. This suggests that there is ~20% in the reads separating the reference replicate from the target replicate.

Figure S3 - Illustration of the noise estimation method comparing a reference sample to a technical replicate as noise is simulated

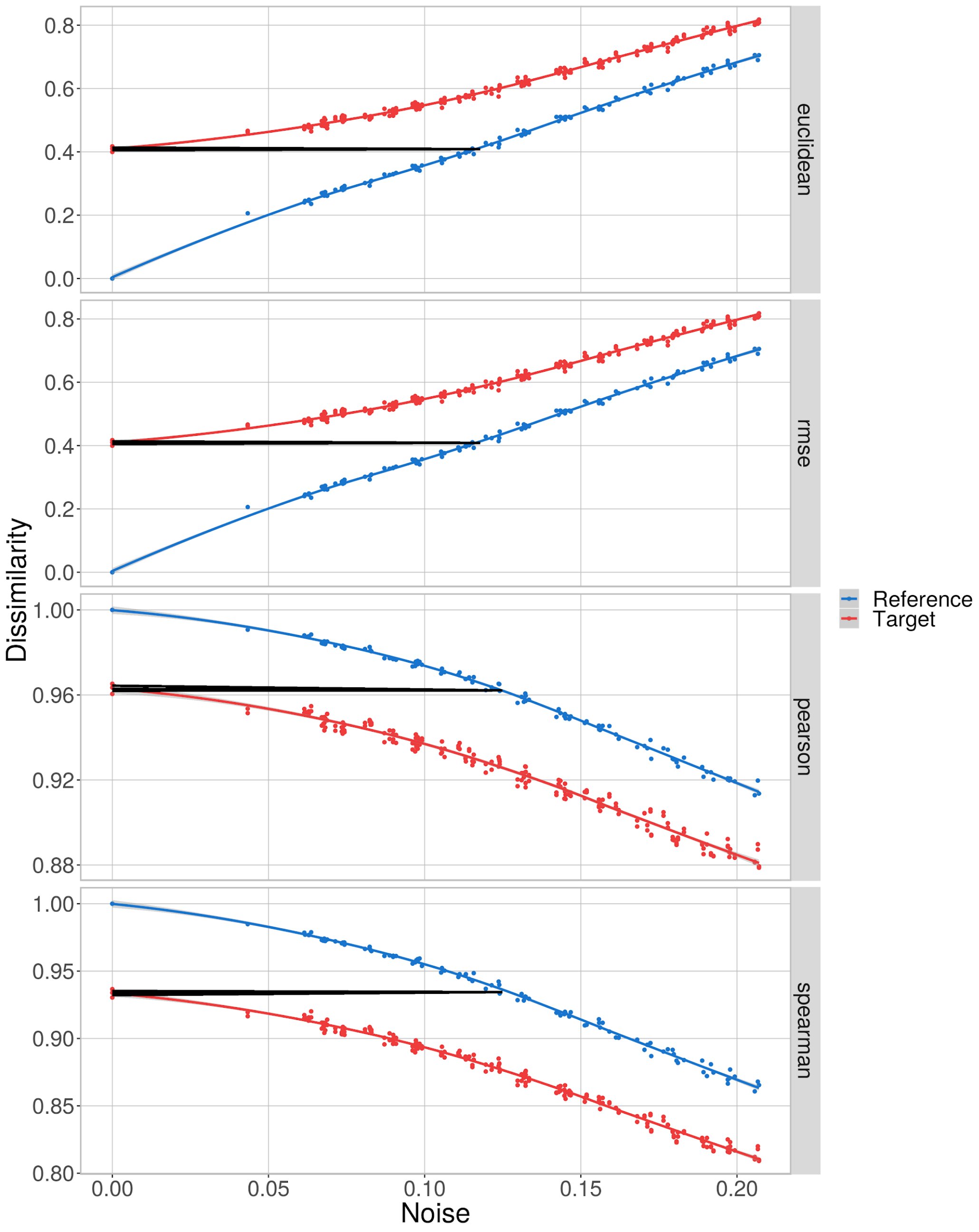


Simulated noise trajectories for reference (blue)--underwent one freeze-thaw--and corresponding target replicate (red)--underwent two freeze-thaws--of patient 1985. From top to bottom, noise is estimated using Euclidean distance, RMSE, Pearson correlation, and Spearman correlation. The

x-axis value corresponding to the intersection of the black and blue line approximates 12% noise between the target and reference

### Figure S4 - Illustration of the method used to generate the subsets used for differential expression analyses

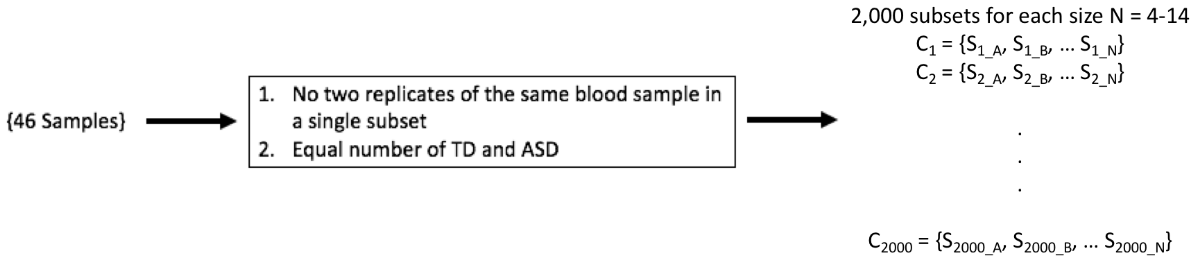


Workflow of bootstrapping procedure for a given subset size N. The total number of samples, including replicates, is 46. Requirements (boxed) are applied to the subsetting, generating 2000 random subsets for each size N. In detail, each DE iteration underwent two subsetting steps. In the first, 2000 subsets of size 16 were generated that met requirement #1. Requirement #1 enforced a maximal size of 16 to the subsets. In some cases, the maximal size could only be 14 due to one of the replicates not having a measured RIN value. Thus, in the second round of subsetting, each maximal size subset was further subsetted to generate 2 subsets per subset size ranging between 4 and 14 under requirement #2.

Figure S5 - Aggregate MA plots for subset differential expression analysis

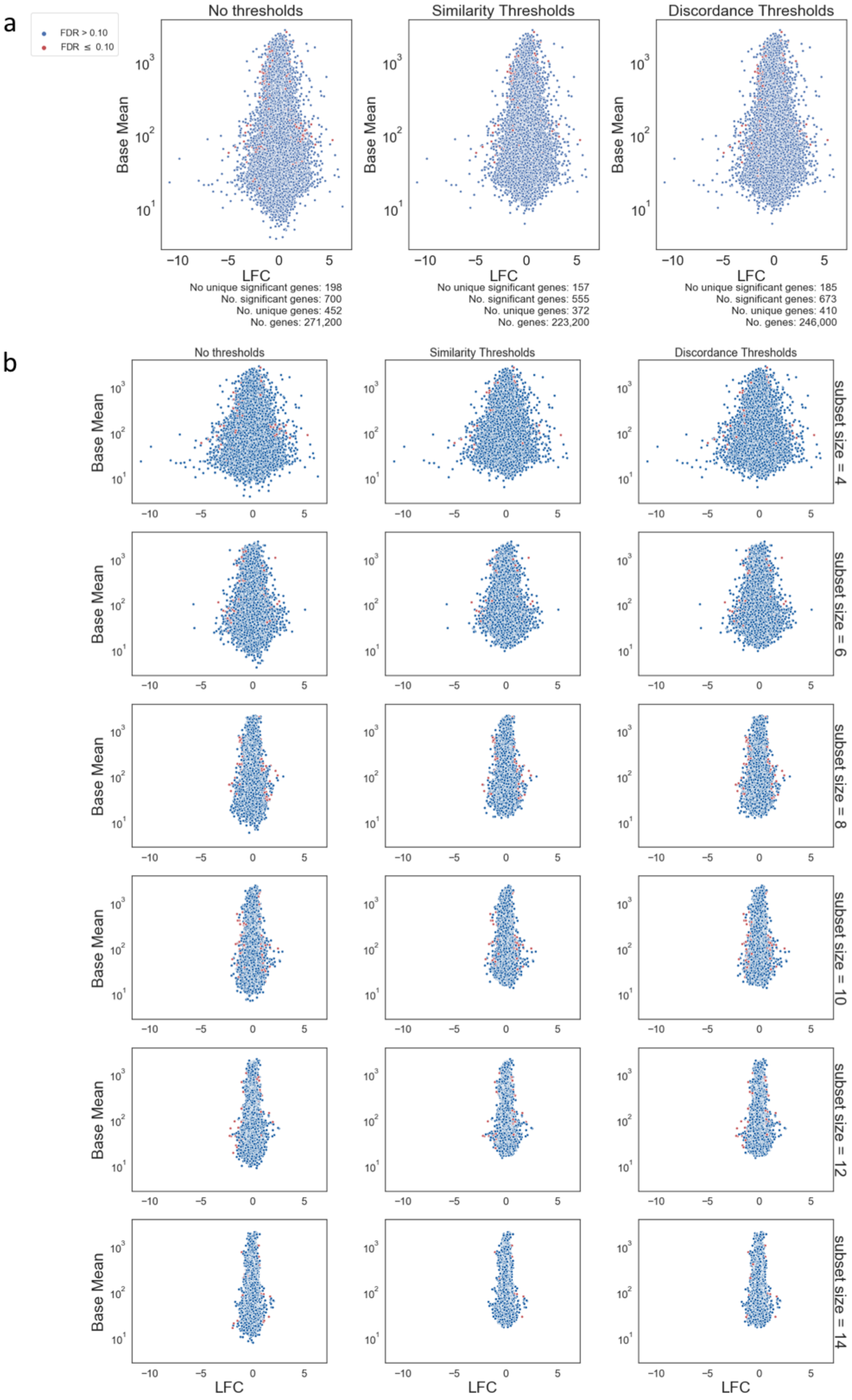
a
FDR > 0.10
FDR ≤ 0.10
No thresholds
Similarity Thresholds
Discordance Thresholds
Base Mean
LFC
No unique significant genes: 198
No. significant genes: 700
No. unique genes: 452
No. genes: 271,200
No unique significant genes: 157
No. significant genes: 555
No. unique genes: 372
No. genes: 223,200
No unique significant genes: 185
No. significant genes: 673
No. unique genes: 410
No. genes: 246,000
b
subset size = 4
subset size = 6
subset size = 8
subset size = 10
subset size = 12
subset size = 14

MA plots comparing base mean (y-axis) to LFC (x-axis) from DE results run on subsets. 5% of all subsets and 10% of all genes were randomly sampled before generating the plots. Genes that are identified as differentially expressed (FDR < 0.1) are colored red. From left to right, panels display genes without any thresholding, the genes remaining after thresholding for similarity, and the genes remaining after thresholding for discordance. (A) MA plots considering all subsets. Labels for number of genes considers total genes across all subsets, whereas labels for number of unique genes only considers non-redundant genes across all subsets. (B) MA plots for genes stratified by subset size, with N = 4 in the top row and N = 14 in the bottom row.

### Figure S6 - Illustration of similarity calculation between subset differential expression analyses

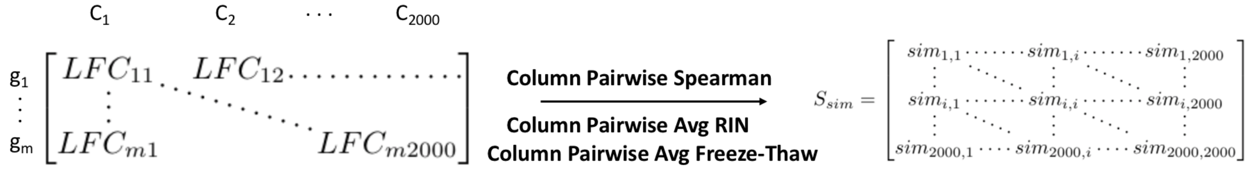


A demonstration of how the similarity scores were calculated for all subsets of a given size N from DE log-fold change results. The pairwise spearman correlation between subsets was measured from the gene ($g_i$) by subset ($C_j$) log-fold change matrix. The result was a symmetric matrix of similarity scores ($S_{sim}$), each element of which represented a subset pair. Additionally, for each subset pair, the average RIN and average freeze-thaw across all samples in the pair was calculated.

### Figure S7 - Similarity, RIN, and freeze-thaw distributions in similarity analyses

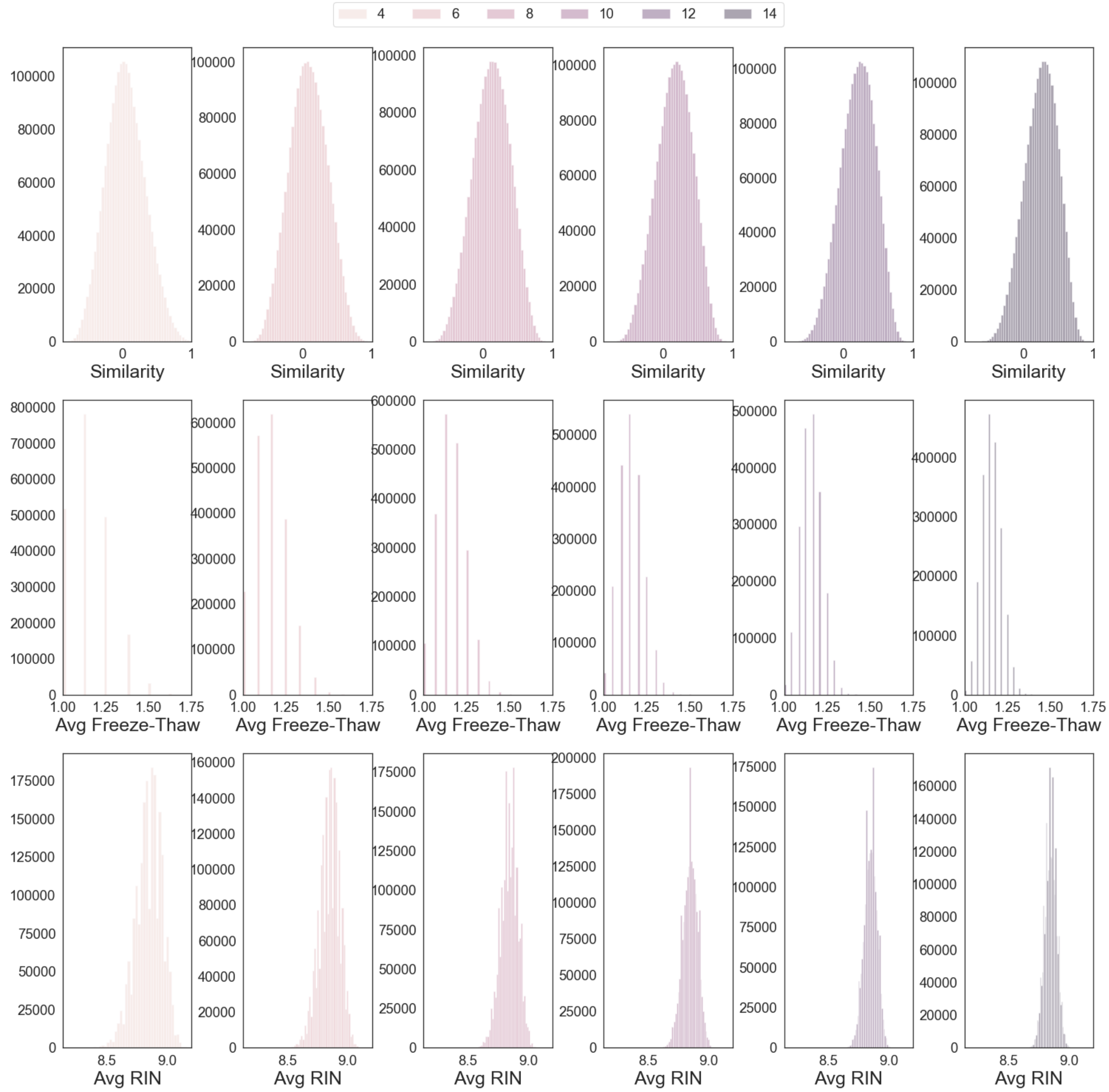


Histograms for the distribution of various metrics in similarity analyses for DE reproducibility. From top to bottom, metrics include similarity (top), Average Freeze-Thaw Cycles (middle) , and Average RIN (bottom) for compared subset pairs. Panels from left to right display distributions for subsets of increasing size (N = 4 - 14).

### Figure S8 - RIN and freeze-thaw distributions across various subset sizes for discordance analysis

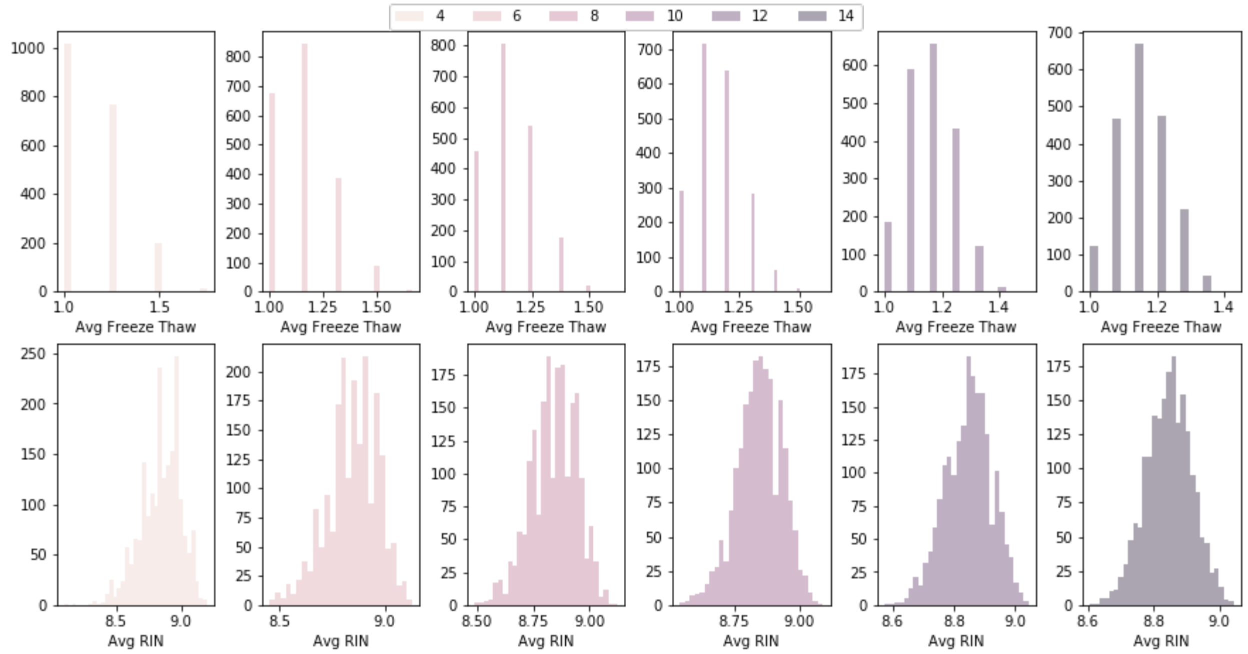


Histograms for the distribution of various metrics in discordance analyses for DE reproducibility. Metrics include Average Freeze-Thaw Cycles (top) and Average RIN (bottom) for each subset. Panels from left to right display distributions for subsets of increasing size (N = 4 - 14).

### Figure S9 - Method for calculation of discordance

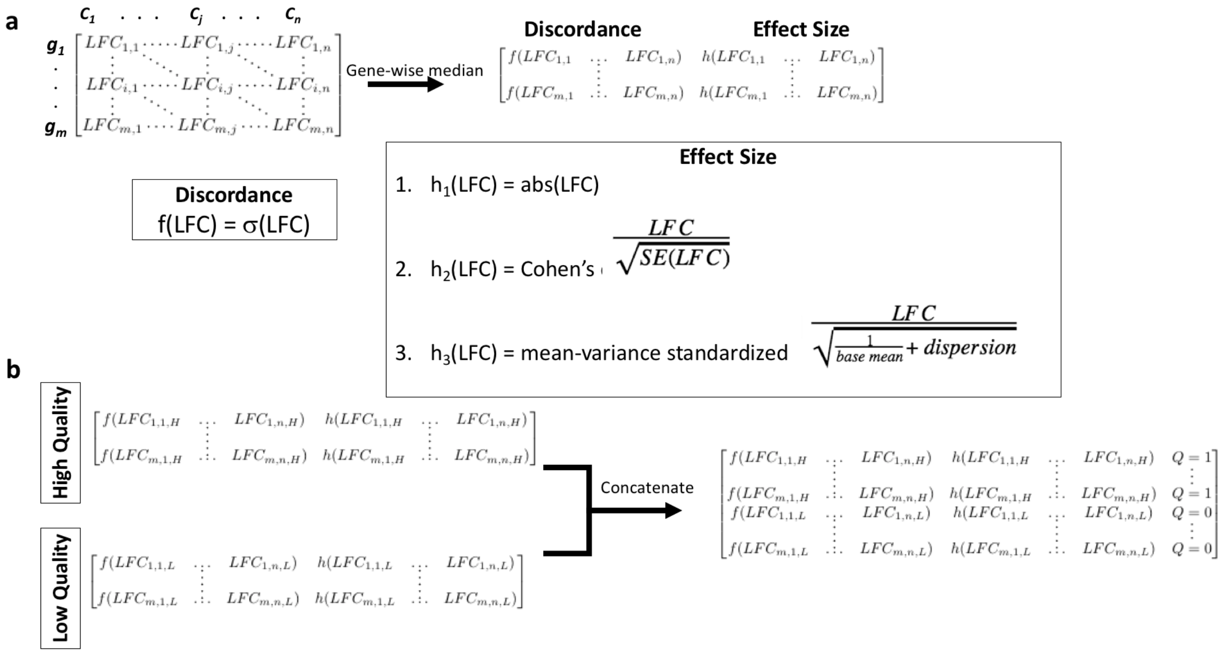


Method for calculating discordance. (A) Subsets ($C_j$) were selected either by subset size or by subset size and sample quality (freeze-thaw or RIN) bin to generate the gene ($g_i$) by subset matrix. The standard deviation was calculated as discordance. This was compared to the gene-wise median effect size as measured by the absolute value of the median LFC, Cohen's d, or mean-variance standardized effect. Each variable in the respective equations $h_1$-$h_3$ is first calculated by the gene-wise median. (B) For GLM(Discordance~Effect Size x Sample Quality), at each subset size, discordance and effect size for high and low quality subsets was concatenated with the addition of a dummy variable Q. Q = 1 for high quality subsets and 0 for low quality subsets.

#### Figure S10 - Distribution of differential expression similarities for different subset sizes

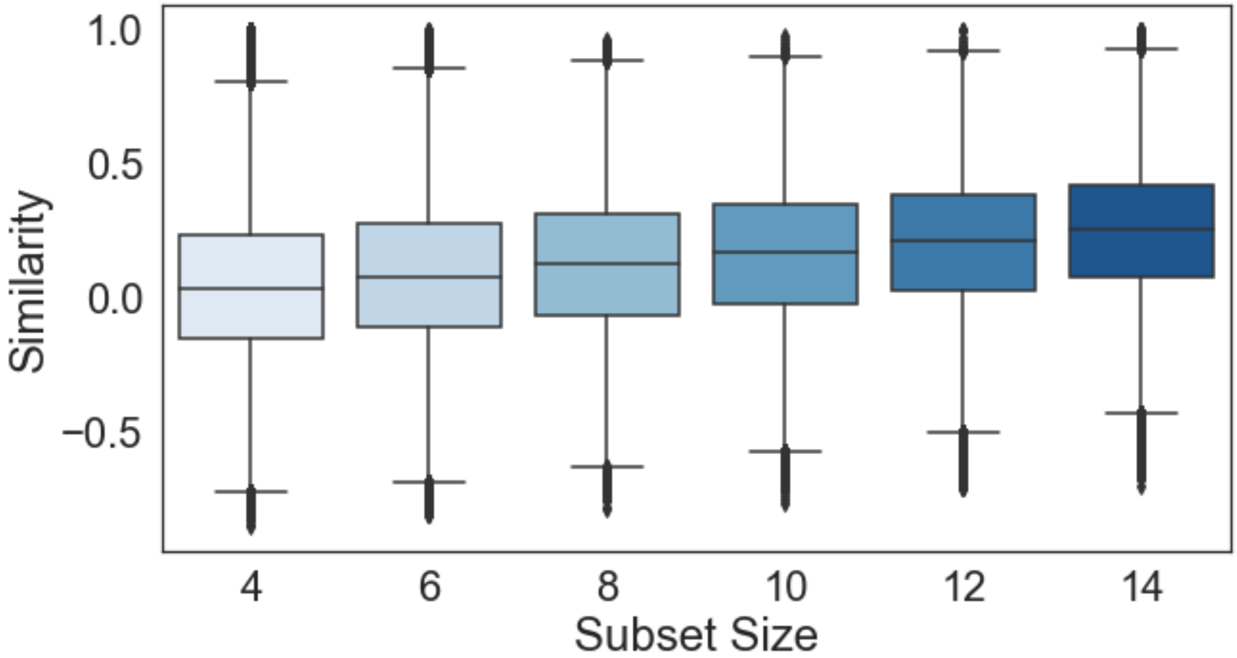


Box plots of similarity, correlation, at each subset size.

Figure S11 - Differential expression discordance for different subset sizes

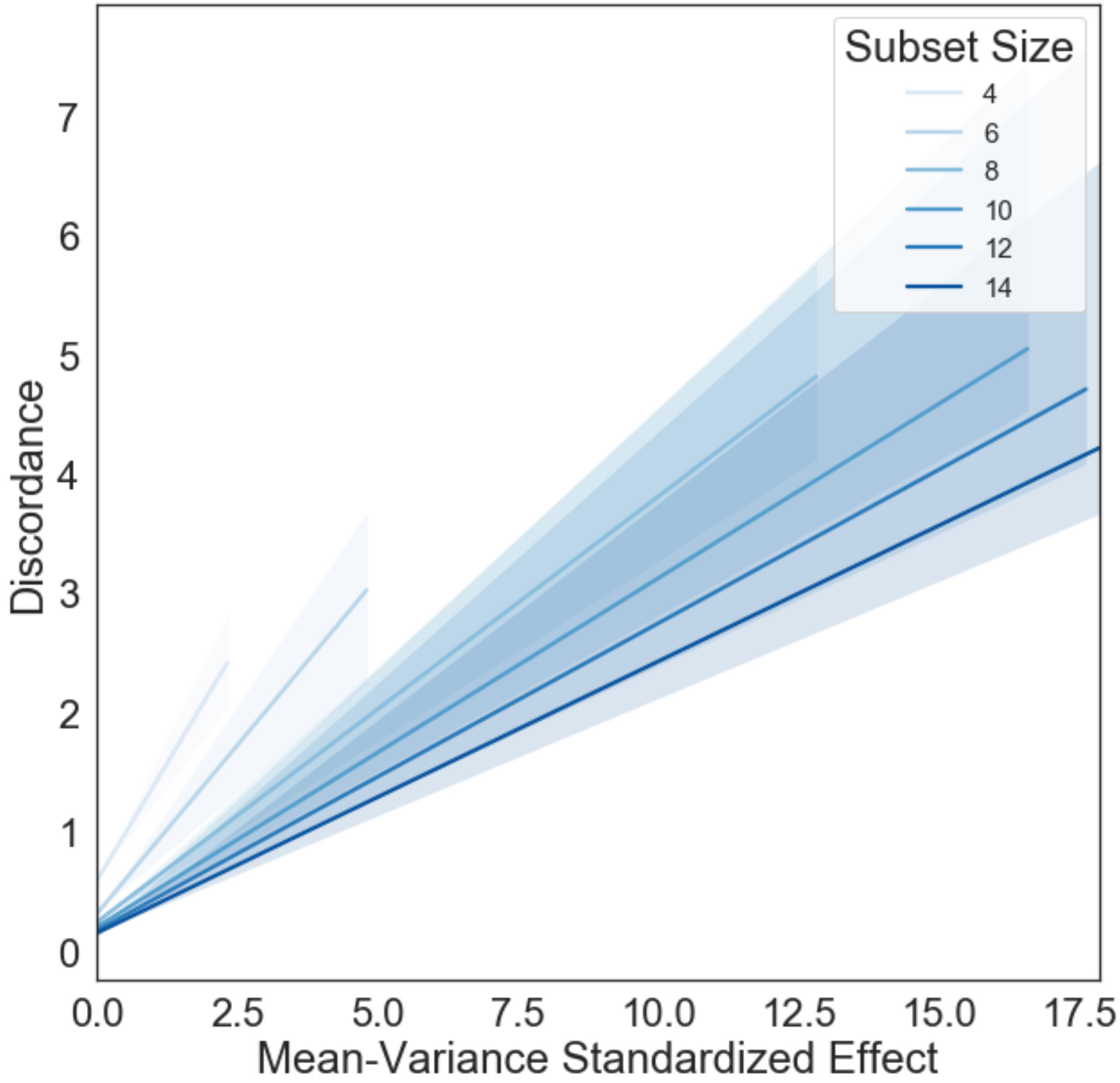


Linear regression estimates for discordance (y-axis) changing as a function of gene-wise effect size measured as mean-variance standardized effect (x-axis). Shaded regions represent 95% confidence intervals for each regression.

### Figure S12 - Concentration does not affect noise

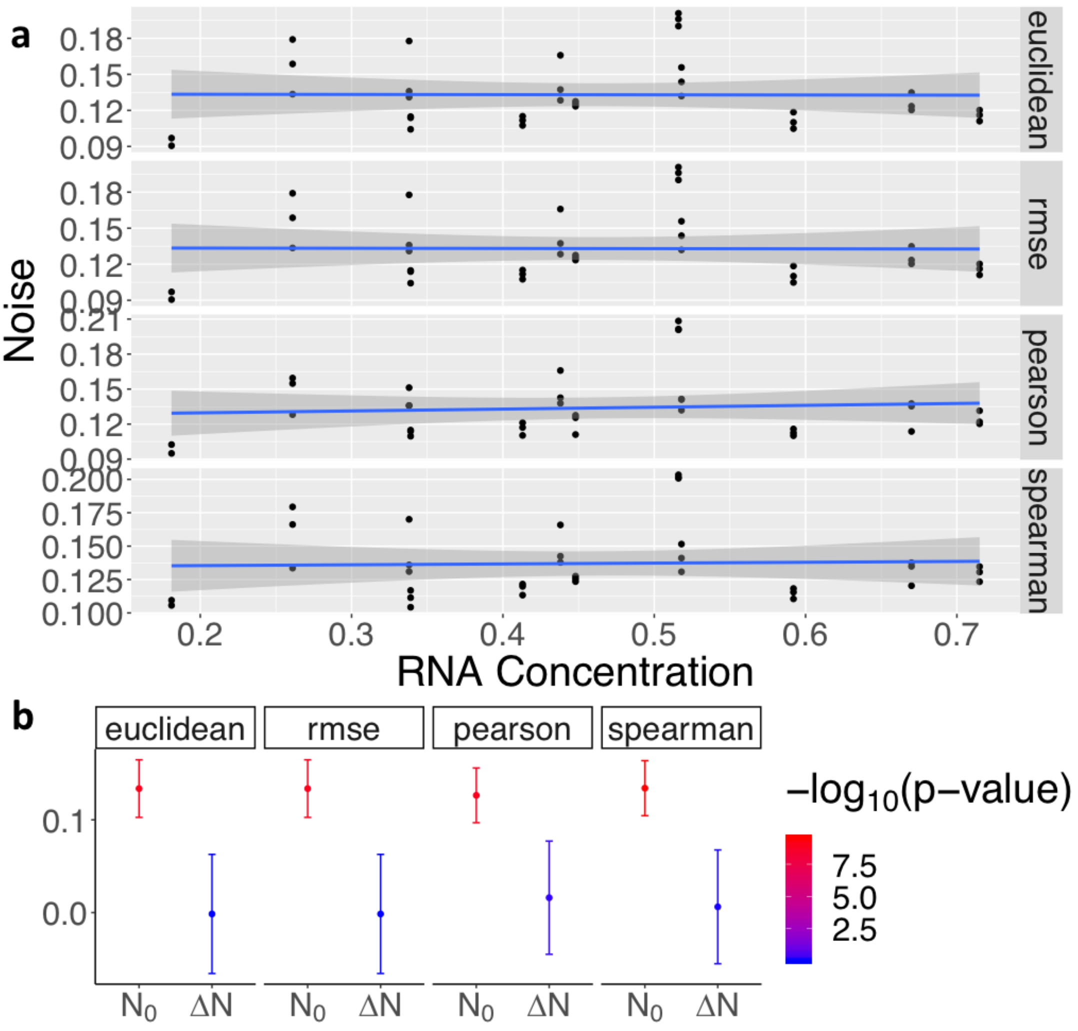


Noise as calculated by Euclidean distance, RMSE, Pearson correlation, and Spearman correlation. (A) Scatter plots comparing noise (y-axis) to RNA concentration [ng/uL] (x-axis).The blue line shows the regression fit and the shaded area is the 95% confidence interval for this fit. (B) GLM (Noise ~ RNA Concentration) intercept ($N_0$) and coefficient values ($\Delta N$) with standard error bars and p-values colored in -log10 scale for each metric.

### Figure S13 - 3’ Bias in polyA-extracted samples

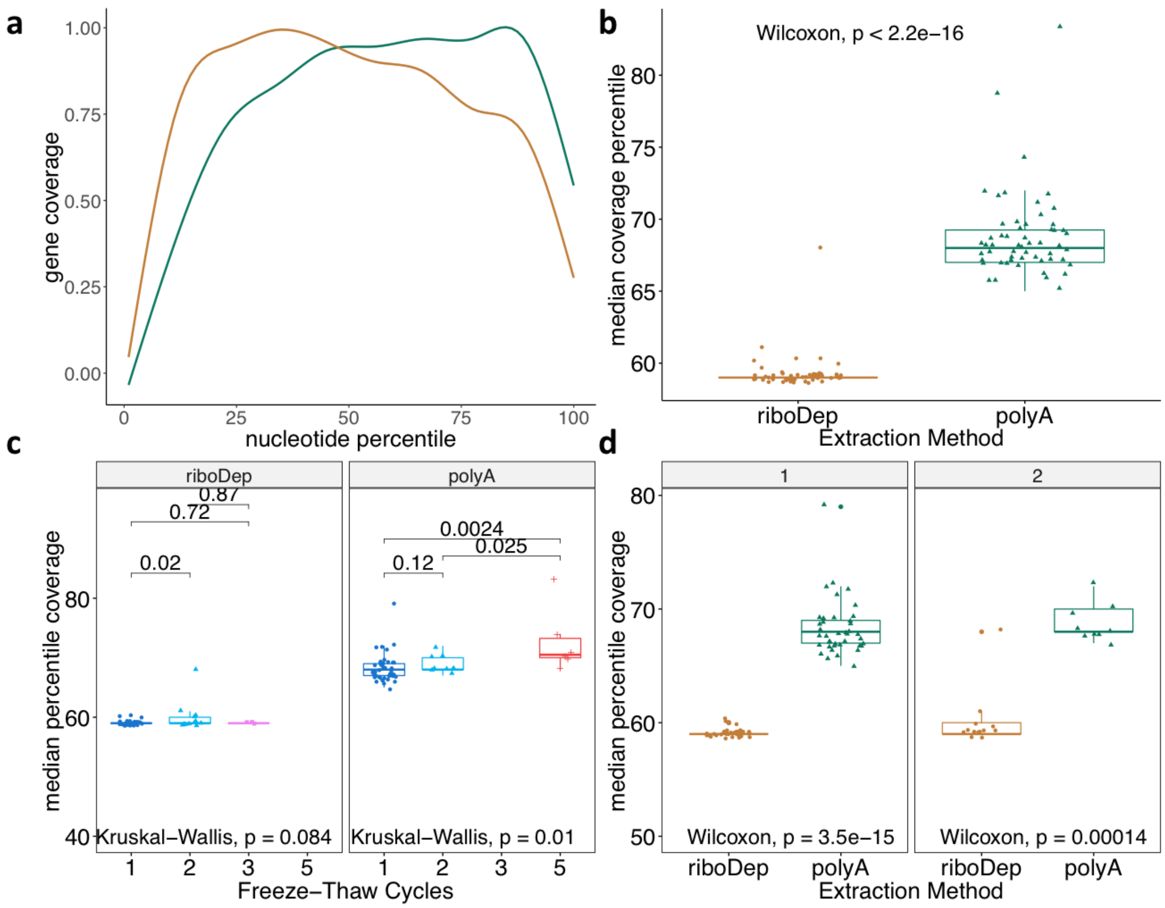


Comparisons of coverage for samples that underwent poly(A)-enrichment (green) or ribosome depletion (orange). (a) Gene coverage (y-axis) at the $i^{th}$ nucleotide percentile (x-axis). For each sample, coverage is averaged across all genes; samples are aggregated using generalized additive model smoothing. (b) Boxplots comparing median coverage percentile for each enrichment method. (c) For each enrichment method, boxplots comparing median coverage percentile between samples that underwent 1-5 freeze-thaws. Resultant p-values using a one-sided

Wilcoxon test comparing each sample are displayed above the boxplots. (d) Box plots comparing median nucleotide position between enrichment methods for either one (left) or two (right) freeze-thaw cycles.

#### Figure S14 - Distribution of RIN and concentration with respect to noise

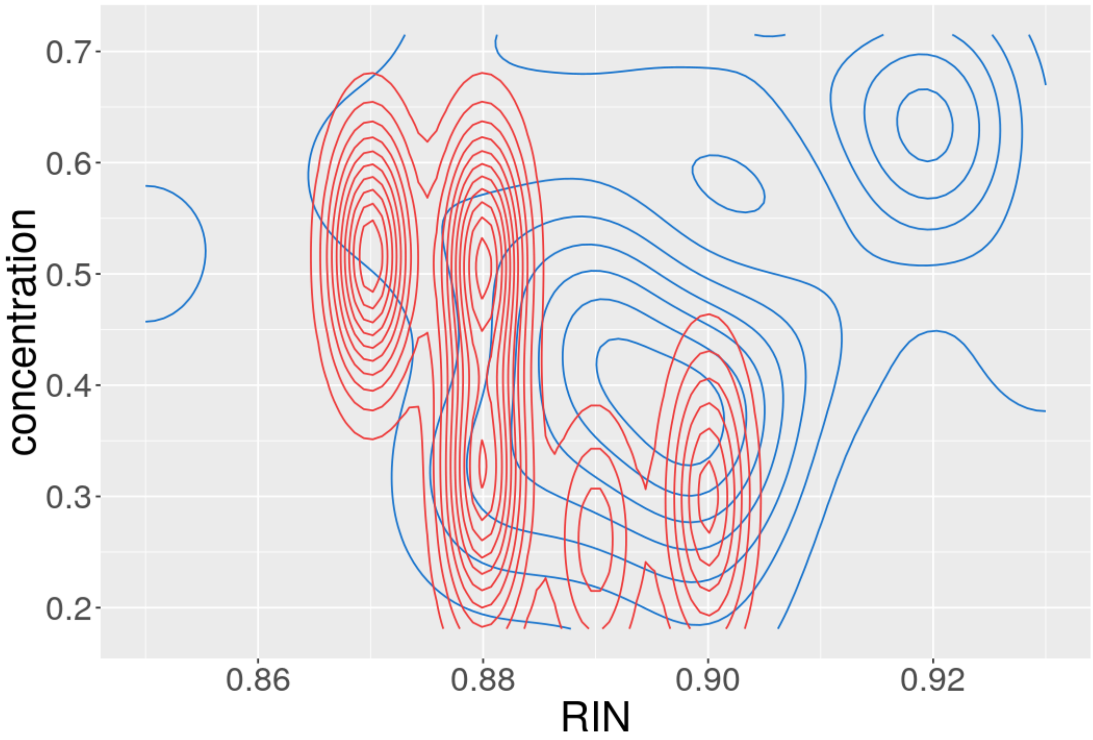


Two dimensional distribution of samples exposed to one freeze-thaw cycles with respect to RIN (x-axis) and concentration (y-axis). High noise (noise > 0.17) samples (red) tend to have lower RIN and concentration than normally performing samples (blue). This distribution is particularly skewed for RIN between high noise and normally performing samples.

#### Figure S15 - Projected random counts for as a function of library size, number of active genes, and percent noise

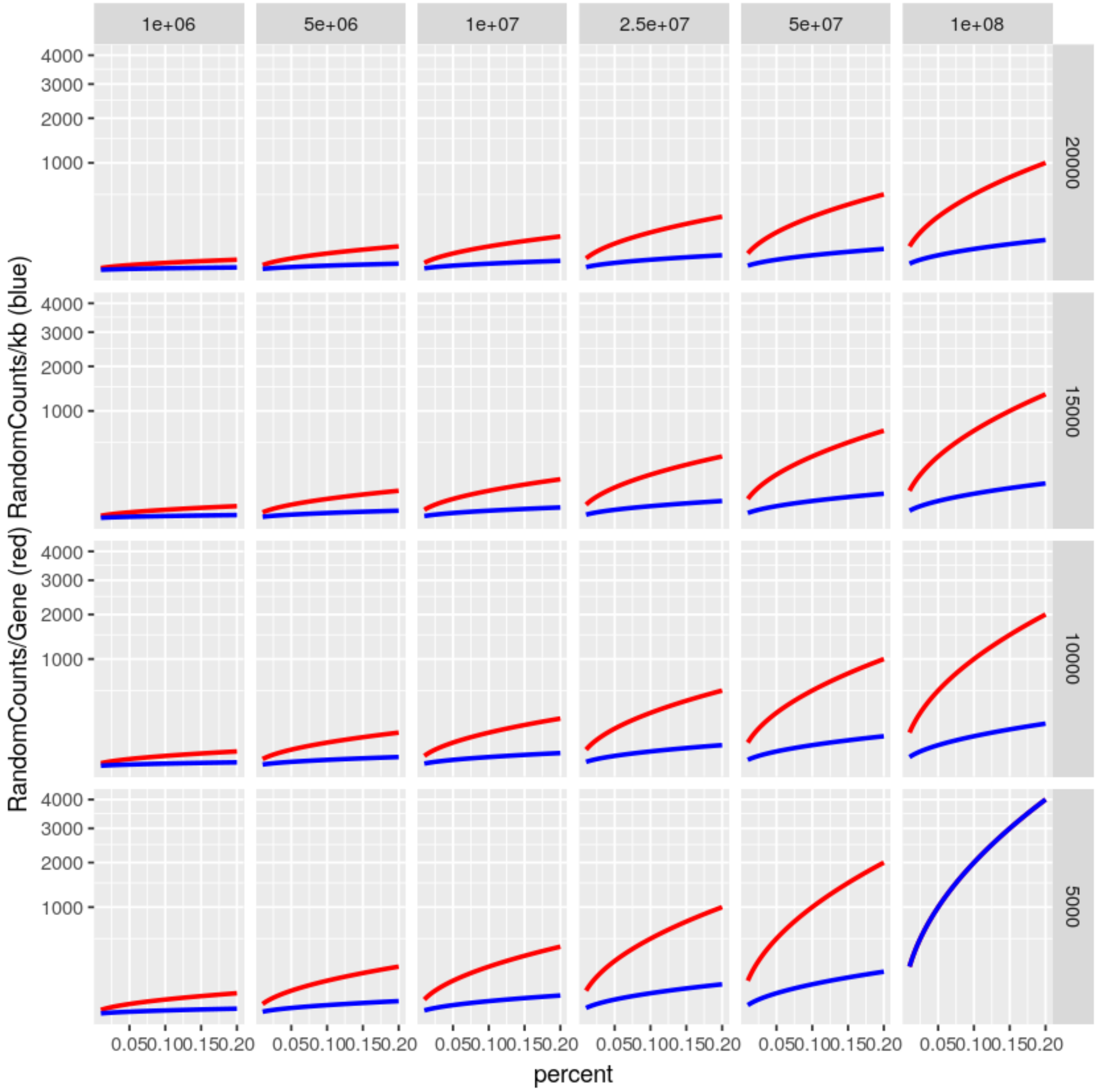


Expected behavior for random counts per gene (red) and kilobase (blue) given the percent noise measured in a sample; the proportion of random counts in a sample as measured by consistency between technical replicates. Randomness is stratified given the number of active genes in the genome (5,000 - 20,000) and the library size (1M to 100M reads); both metrics assume uniform expression to estimate a median random count value. Random counts per gene assumes a constant gene size and random counts per kilobase assumes a constant average of 12.5 kilobases per gene. The number of random counts increases rapidly as percent noise, number of active genes and library size increase.

### Supplementary Methods

#### Simulation of Technical Replicates

Simulated replicates were generated by iteratively selecting genes from a uniform distribution over the genome. From the selected gene, *n* reads are added or subtracted, where *n* is drawn from a Poisson ($\lambda = C$) and *C* is the number of reads aligned to that gene. Iteration stops once dissimilarity between simulated and reference replicate matches dissimilarity between simulated and target replicate. Using a Poisson distribution to simulate noise between technical replicates is standard [1–4]. The overdispersion of the Poisson distribution into the negative binomial distribution is thought to result from the addition of biological variation to the base technical Poisson variation [5].

The percent noise *n* can be used to calculate the number of random counts per gene (RCPG). Given the total number of protein-coding genes in the genome of interest $N_g$ and a library size $L$:

$$RCPG = n * \frac{L}{N_g}$$

#### Freeze-Thaw GSEA

We were interested in exploring any potential functional impact of sample quality; whether or not sample quality disproportionately impacted any functional gene groups. To do this, we examined differential expression with respect to freeze-thaw. DE between one and two freeze-thaw cycles was conducted using DESeq2 [5]. We filtered our expression matrix for genes with an average count less than or equal to 20 across all samples. This reduced the number of genes from 10,028 to 4,520. We used between zero and four covariates calculated from either SVA[6] or RUV.

For our analysis, we retained DE genes with a base mean > 10, |LFC| > 5, and an adjusted p-value < 0.1. We used each filtered DE result as input to gene set enrichment analysis using the gseGO function from clusterProfiler [7]. We performed enrichment using gene sets with 10-500 genes from the gene ontology (GO) biological processes, a p-value cutoff of 0.05 after 1000 permutations. Finally, we calculated the pairwise Jaccard index of significantly enriched terms between each result to determine the consistency of enrichments.

### Supplementary Results

#### Changes in Concentration Have a Negligible Effect on Noise

As a negative control, we examined the effect of noise due to changes in RNA concentration. We did not expect to see any effect, as concentration within a normal range is not a known source of noise between technical replicates. We observe a negligible effect on noise due to changes in concentration ranging between 20-75 ug/mL. The base noise is comparable to the effect observed in the freeze-thaw cycle examination, suggesting that samples are consistent between the two analyses (**Fig. S12**).

#### Using RUV in DE Recovers an Autism Signal

We used RUV to introduce covariates--latent control variables--to the DE design matrix (**Supplementary Methods**). RUV controls for unknown sources of unwanted variation. We check the ability of RUV to remove unwanted variance, thus revealing DE genes associated with autism. We used a previously published, twice validated [8,9], Autism (ASD) gene expression signature to validate our signature. At a |LFC| threshold ≥ 0.2, RUV-normalized DE showed enrichment ($p$=0.0022, hypergeometric enrichment test) for in the known Autism signature (data not shown).

#### GSEA Does Not Identify an Apparent Freeze-Thaw Gene Signature

We ran GSEA on 9 similar DE analyses across one and two freeze-thaw samples. Of these 9, only 4 identify > 1 enriched terms (**Supplementary Table 8-9**). In other words, only 6 of the total 36 pairwise comparisons between GSEA results could possibly have a Jaccard index > 0. Half of these have a Jaccard index < 0.1. We find a median Jaccard index of 0.05 when disregarding analyses that resulted in ≤ 1 enriched terms.The smallest number of enriched terms when disregarding analyses that results in ≤ 1 enriched term is 15.

### Supplementary Discussion

We note that, although samples with a single freeze-thaw cycle are significantly less noisy than matched samples with two freeze-thaw cycles, there are still high noise samples with only one freeze-thaw (**Fig. 2)**. High noise samples that have only undergone one freeze-thaw are likely to have low RIN or low concentration (**Fig. S14)**.

Given the limited number of enriched terms and the low Jaccard index value found between terms, we infer that there is no clear gene signature to freeze-thaw. In other words, there is not a list of transcripts that are more prone to degradation and other freeze-thaw effects than random.
